# Transcriptomics at maize embryo/endosperm interfaces identify a novel transcriptionally distinct endosperm sub-domain adjacent to the embryo scutellum (EAS)

**DOI:** 10.1101/799338

**Authors:** Nicolas M. Doll, Jeremy Just, Véronique Brunaud, José Caïus, Aurélie Grimault, Nathalie Depège-Fargeix, Eddi Esteban, Asher Pasha, Nicholas J. Provart, Gwyneth C. Ingram, Peter M. Rogowsky, Thomas Widiez

**Affiliations:** Laboratoire Reproduction et Développement des Plantes, Univ Lyon, ENS de Lyon, UCB Lyon 1, CNRS, INRA, F-69342, Lyon, France; Institute of Plant Sciences Paris Saclay IPS2, CNRS, INRA, Université Paris-Sud, Université Evry, Université Paris-Saclay, Bâtiment 630, F-91405 Orsay, France; Institute of Plant Sciences Paris-Saclay IPS2, Paris Diderot, Sorbonne Paris-Cité, Bâtiment 630, F-91405, Orsay, France; Department of Cell and Systems Biology/Centre for the Analysis of Genome Evolution and Function, University of Toronto, Toronto, Ontario M5S 3B2, Canada

## Abstract

Seeds are complex biological systems comprising three genetically distinct tissues nested one inside another (embryo, endosperm and maternal tissues). However, the complexity of the kernel makes it difficult to understand inter compartment interactions without access to spatially accurate information. Here we took advantage of the large size of the maize kernel to characterize genome-wide expression profiles of tissues at embryo/endosperm interfaces. Our analysis identifies specific transcriptomic signatures in two interface tissues compared to whole seed compartments: The scutellar aleurone layer (SAL), and the newly named endosperm adjacent to scutellum (EAS). The EAS, which appears around 9 days after pollination and persists for around 11 days, is confined to one to three endosperm cell layers adjacent to the embryonic scutellum. Its transcriptome is enriched in genes encoding transporters. The absence of the embryo in an *embryo specific* (*emb*) mutant can alter the expression pattern of EAS marker genes. The detection of cell death in some EAS cells together with an accumulation of crushed cell walls suggests that the EAS is a dynamic zone from which cell layers in contact with the embryo are regularly eliminated, and to which additional endosperm cells are recruited as the embryo grows.

## Introduction

Cereal grains are not only essential for plant propagation, but are also high value products which represent an important source of calories and proteins for human nutrition and animal feed, as well as a coveted resource for bio-sourced industries. In maize, the accumulation of oil in the embryo and of starch and protein in the endosperm requires the development of adapted structures and a coordinated regulation and distribution of the nutrient flow from the mother plant. The development of the embryo, which will form the future plant, and the endosperm, which will nourish the embryo during germination occurs in three main phases (Berger, 1999; Dumas and Rogowsky, 2008; Lopes and Larkins, 1993). During the first two weeks of early maize seed development, embryo and endosperm cells differentiate into populations forming distinct tissues and organs (Leroux et al., 2014; Randolph, 1936), including two storage organs, the scutellum of the embryo and the starchy endosperm (*early development phase*). These two zygotic compartments then start to accumulate large quantities of storage compounds during the following two to three weeks (*filling phase*), while the surrounding maternal tissues provide or transport the necessary nutrient supplies (Porter et al., 1987; Wu and Messing, 2014). During the final four weeks (*maturation phase*), the kernel dehydrats and enters into quiescence prior to dispersal (Sabelli and Larkins, 2009; Sreenivasulu and Wobus, 2013; Vernoud et al., 2005). These three phases are determined by distinct genetic programs and characterized by distinct anatomical and cytological features. Spatially, the maize kernel is organized like Russian dolls, the embryo being enclosed within the endosperm, which is itself surrounded by the pericarp.

A closer look at the highly differentiated structure displayed by the maize embryo shows that four days after pollination (DAP) two distinct parts can be distinguished: an apical embryo proper and a basal suspensor that will degenerate at the end of early development (Doll et al., 2017; Giuliani et al., 2002). At around 8 DAP the embryo proper generates, at the abaxial side, a shield-shaped organ, the above-mentioned scutellum, and the shoot apical meristem at the adaxial side. Marking the apical pole of the future embryonic axis, the shoot apical meristem will produce several embryonic leaves over time. The root apical meristem differentiates within the embryo body defining the basal pole of the embryonic axis. Shoot and root meristems will respectively surrounded by the protective coleoptile and coleorhiza (Bommert and Werr, 2001; Randolph, 1936; Vernoud et al., 2005). The surrounding endosperm, which occupies 70% of the kernel volume at the end of the early development (Leroux et al., 2014; Rousseau et al., 2015; Sabelli and Larkins, 2009; Zhan et al., 2017), has been described as differentiating only four main cell types. The basal endosperm transfer layer (BETL) and the aleurone layer (AL) are two peripheral cell types in contact with maternal tissues. The embryo-surrounding region (ESR) is formed of small densely cytoplasmic cells encircling the young embryo. Lastly, the starchy endosperm (SE) corresponds to the central region of the endosperm, which subsequently accumulates huge amounts of storage compounds before undergoing progressive programmed cell death. The developing endosperm is surrounded by maternal tissues: the nutritive nucellus that degenerates as the endosperm expands, and the protective pericarp, which comprises the pedicel at the basal pole (Berger, 2003; Olsen, 2001; Sabelli and Larkins, 2009; Zhan et al., 2017).

The parallel growth and profound developmental changes of the three embedded kernel compartments highlight the need for constant coordination, which likely requires a complex inter-compartmental dialog (Ingram and Gutierrez-Marcos, 2015; Nowack et al., 2010; Widiez et al., 2017). Since maternal tissues, endosperm and embryo are symplastically isolated, their apoplastic interfaces represent essential zones for this dialog (Diboll and Larson, 1966; Van Lammeren, 1987). A good example to illustrate the importance and specialisation of interfaces is carbon transport. Sugars have to be transported from the maternal tissues to the embryo for growth and fatty acid accumulation, passing through the endosperm, which needs to retain part of the carbon for its own growth as well as the biosynthesis of starch and storage proteins (Chourey and Hueros, 2017; Sabelli and Larkins, 2009). In maize, nutrients are unloaded from open ends of the phloem vessels into the placento-chalazal zone of the maternal pedicel (Bezrutczyk et al., 2018; Porter et al., 1987). At the base of the endosperm, the BETL cells form dramatic cell wall ingrowths, thus increasing the exchange surface (Davis et al., 1990; Kiesselbach and Walker, 1952). BETL cells express a specific set of genes, including *MINIATURE1*, encoding a cell wall invertase, which cleaves sucrose into hexoses. These are taken up by the sugar transporter SWEET4c, which has been demonstrated to be the key transporter of sugar at the pedicel/endosperm interface, since the defects in seed filling of the corresponding *sweet4c* mutant lead to a miniature kernel phenotype (Cheng et al., 1996; Kang et al., 2009; Lowe and Nelson, 1946; Sosso et al., 2015). The remaining endosperm interface with maternal tissues (initially the nucellus and later on the pericarp) is the AL, which is not known to contribute to nutrient exchange during seed development (Gontarek and Becraft, 2017).

The interface between the endosperm and the embryo is also developmentally dynamic. At 3-6 DAP, the embryo is totally surrounded by ESR-type cells. As the embryo expands, it emerges from the ESR, which consequently becomes restricted to the zone surrounding the basal part (suspensor) of the embryo and ultimately disappears together with the suspensor at the end of the early development phase (Giuliani et al., 2002; OpsahlFerstad et al., 1997). From 8-9 DAP, the upper part (embryo proper) forms two new interfaces: (1) At the adaxial side the embryo is enclosed by a single cell layer, which is called the scutellar aleurone layer (SAL) in barley (Jestin et al., 2008). (2) At the abaxial side, the embryo is brought into direct contact with central starchy endosperm cells (Van Lammeren, 1987). This interface is constantly moving due to the growth of the scutellum inside the endosperm. On the embryo side of this interface, the epidermis of the scutellum has a distinct morphology and gene expression pattern (Bommert and Werr, 2001; Ingram et al., 2000). The dynamics of the endosperm/embryo interface, and the processes that occur there, remain largely undescribed.

At many inter-compartmental interfaces, such as the BETL, the ESR or the AL, the cells constitute readily identifiable tissues with distinctive and often striking cell morphologies, with defined organisations and established functions (except for the ESR) (for review see Doll et al., 2017). In many cases specific sets of genes are expressed in these tissues, as revealed by the identification and characterisation of marker genes, for example of *MATERNALLY EXPRESSED GENE 1* (*MEG1)*, *MYB-RELATED PROTEIN 1 (MRP1) and BETL1* to *4* (Cai et al., 2002; Gómez et al., 2002; Gutiérrez-Marcos et al., 2004; Hueros et al., 1999a, 1999b) in the BETL *, VIVIPARIOUS 1 (VP1)* in the AL (Suzuki et al., 2003), or *ESR1* to *3* in the ESR (OpsahlFerstad et al., 1997).

Genome-wide gene expression studies at numerous developmental stages of whole kernels and/or hand-dissected endosperm and embryo (Chen et al., 2014; Downs et al., 2013; Li et al., 2014; Lu et al., 2013; Meng et al., 2018; Qu et al., 2016) have been complemented by a recent transcriptomic analysis of laser-capture micro-dissected cell types and sub-compartments of 8 DAP kernels (Zhan et al., 2015). However, even the latter study did not address specifically the transcriptomic profiles of the embryo/endosperm interfaces and did not answer the question of whether the endosperm at the scutellum/endosperm interface is composed of cells with specific transcriptional identities.

In this study, we took advantage of the large size of the maize kernel to characterize the genome-wide gene expression profile at embryo/endosperm interfaces at 13 DAP. RNA-seq profiling revealed that endosperm cells in close contact with the embryo scutellum have a distinct transcriptional signature allowing us to define a new endosperm zone named EAS for <u>E</u>ndosperm <u>A</u>djacent to <u>S</u>cutellum, which is specialized in nutrient transport based on GO enrichment analysis. *In situ* hybridization spatially confines the EAS to one to three endosperm cell layers adjacent to the scutellum, whereas kinetic analyses show that EAS is present when the scutellum emerges at around 9 DAP and persists throughout embryo growth, up to approximately 20 DAP. The detection of cell death in EAS together with impaired expression of EAS marker genes in an *embryo specific* mutant suggests that the EAS is a developmentally dynamic interface influences by the presence of neighbouring growing embryo.

## Results

### RNA-seq profiling of 13 DAP maize kernel compartments and embryo/endosperm interfaces

To obtain the gene expression patterns of embryo/endosperm interfaces in maize kernels, six (sub)compartments were hand-dissected for transcriptomic analysis (Figure 1). The three whole compartments were: the maternal tissues excluding the pedicel which were labelled pericarp (Per), the whole endosperm (End), and the whole embryo (Emb) (Figure 1). The sub-compartments corresponding to three distinct embryo/endosperm interfaces were the scutellar aleurone layer (SAL) (the single endosperm cell layer at the adaxial side of the embryo), the apical scutellum (AS) (corresponding to the embryo tip composed uniquely of scutellum tissues without the embryo axis), and a new region that we named endosperm adjacent to scutellum (EAS) corresponding to several layers of endosperm cells in close contact with the scutellum at the abaxial side of the embryo (Figure 1). The tissues were collected from kernels of inbred line B73 (used to establish the maize reference genome) at 13 DAP (embryo size of about 2.5 mm), the earliest developmental stage at which hand dissection of these embryo/endosperm interfaces was feasible, and also the transition from early development to filling phase. For each of the six samples, four biological replicates, each composed of a pool of dissected tissues from two different plants, were produced (Supplemental Table 1). A total of 24 RNA-seq libraries were constructed and sequences in paired-end mode using Illumina HiSeq2000 technology. The resulting reads (on average 52 million reads per sample) were quality checked, cleaned and mapped to the current version of the B73 maize reference genome (AGPv4). On average 95.8% (± 0.4%) of the reads were mapped (Supplemental Figure 1), and of these on average 78.3% (± 5.3%) corresponded to annotated genes. Reads that mapped to multiple genes (10.2% ± 5.3%) or to no gene (5.2% ± 1.1%), as well as ambiguous hits (1.5% ± 0.6%) were filtered (Supplemental Figure 1). A gene was considered to be not expressed if it gives rise to less than 1 read per million. At least 25 000 genes were found to be expressed per replicate, with the largest number found in the SAL (∼30 000 genes expressed, Supplemental Figure 1B). The results generated for each replicate are available in Supplemental Data Set 1.

**Figure 1.**
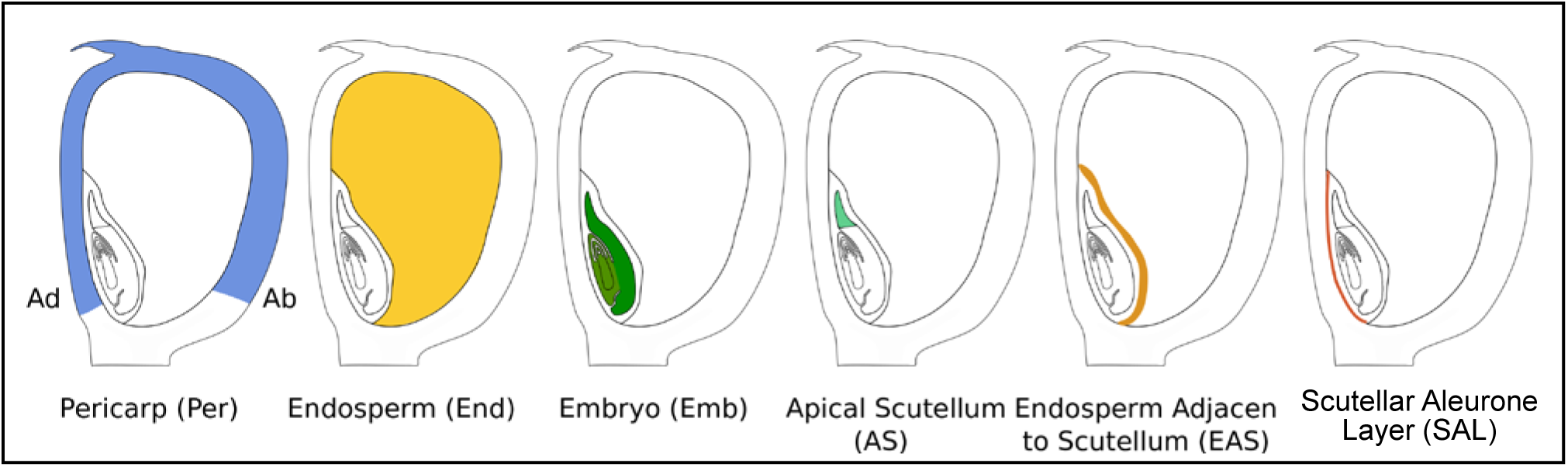
Scheme representing the six (sub)compartments hand-dissected for transcriptomics analysis at maize embryo/endosperm interfaces. Ad = adaxial, Ab = abaxial.

In order to assess the relationships between the different samples, a principal component analysis (PCA) was performed (Figure 2A). As expected, biological replicates grouped together, indicating experimental reproducibility. The PCA also revealed distinct sample populations corresponding to each (sub)compartment with the exception of the AS and Emb samples, which were partially superimposed (Figure 2A). Interestingly, the two endosperm interfaces SAL and EAS formed groups that were distinct both from each other, and from the whole endosperm samples. The EAS was more similar to the whole endosperm than the SAL, indicating a more similar transcriptomic landscape (Figure 2A). To explore potential contamination between tissues during the dissection process, the expression profiles of previously identified marker genes with tissue-specific expression patterns were investigated (Figure 2B-D). *ZmLEC1* (*Zm00001d017898*) and *ZmNAC124* (*Zm00001d046126*, named *ZmNAC6* by Zimmermann and Werr, 2005), two embryo-specific genes, were specifically expressed in the embryo samples in our dataset. As expected, *ZmLEC1* was more strongly expressed in the Emb than in the AS sample (Zhang et al., 2002). Absence of *ZmNAC124* expression in the AS was consistent with the strong and specific *in situ* hybridisation signal for this gene in the basal part of the embryonic axis (Zimmermann and Werr, 2005). The two endosperm-specific genes *ZmZOU*/*O11* (*Zm00001d003677*) and *O2* were found to be strongly expressed in the End and EAS, and weakly in the SAL samples (Figure 2C) (Feng et al., 2018; Grimault et al., 2015; Schmidt et al., 1990). A weak expression in the Per sample was unexpected but consistent with other transcriptomics data (Sekhon et al., 2011), which could also reflect possible contamination of the Per samples by aleurone layer, since the aleurone layer has a tendency to stick to the pericarp (See discussion part). In addition, the preferential expression of *AL9* and *Zm00001d024120* genes in the aleurone was reflected by a stronger signal in SAL compared to End (Gomez et al., 2009; Li et al., 2014; Zhan et al., 2015). Since *AL9* (*Zm00001d012572*) and *Zm00001d024120* also showed a signal in the pericarp samples, indicating again a possible contamination of the Per samples by SAL (Figure 2D).

**Figure 2.**
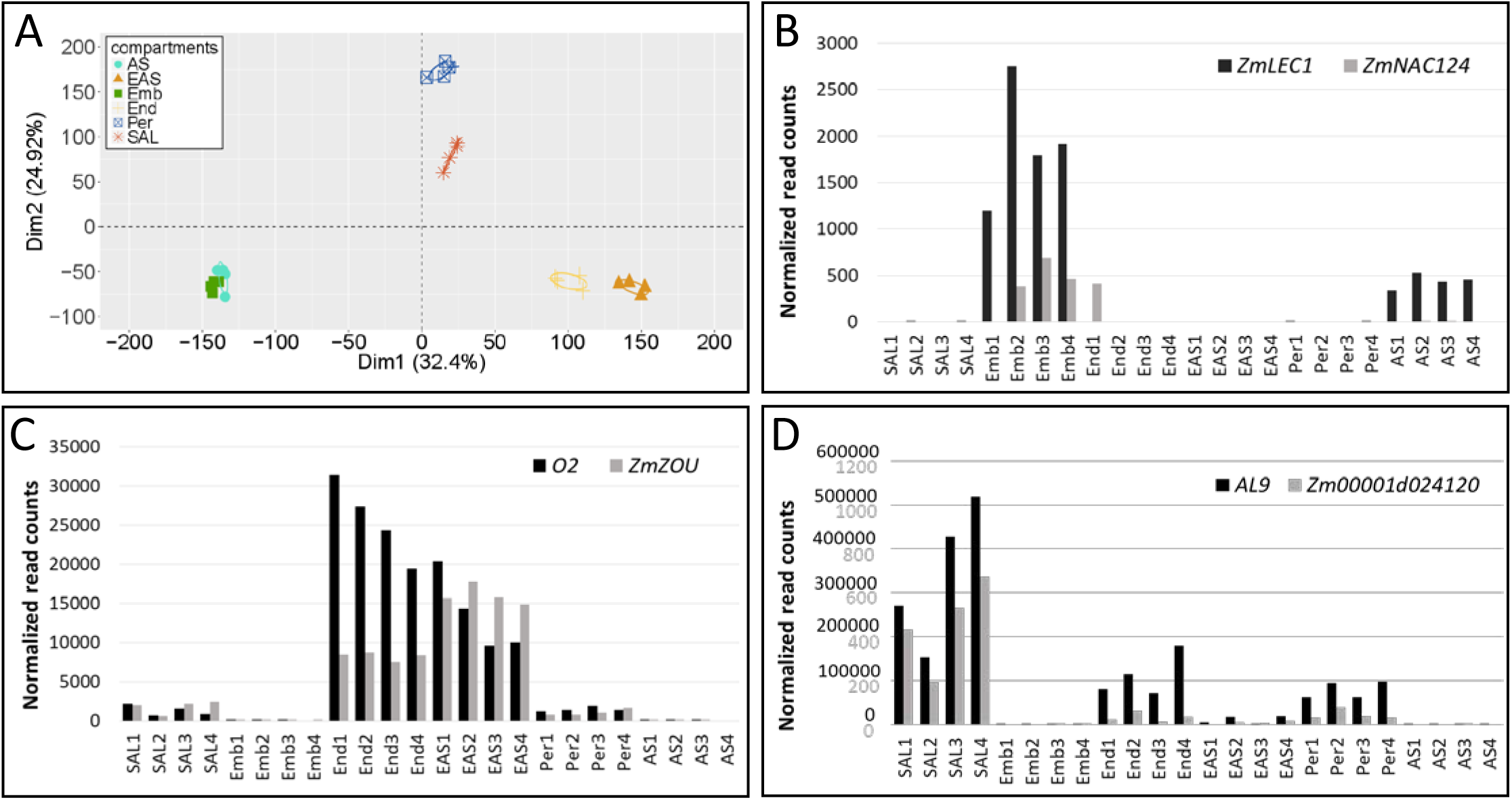
Validation of the RNA-seq approach. **(A)** Principal component analysis of the 24 RNA samples consisting of 4 biological replicates of Pericarp (Per), Apical Scutellum (AS), Embryo (Emb), Endosperm (End), Embryo Adjacent to Scutellum (EAS) and Scutellar Aleurone (SAL). **(B)** to **(D)** graphs represent the expression level (read counts were normalized using the trimmed mean of M-value (TMM) method) in the different samples of **(B)** the two embryo-specific genes LEC1 and ZmNAC124; **(C)** the two endosperm specific genes O2 and ZmZOU (O11) and **(D)** the two aleurone specific genes *AL9* and *Zm00001d024120*. Grey and black Y-scales numbering in **(D)** are for *Zm00001d024120* and *AL9* expression level respectively.

In order to compare our full transcriptomic data set with published RNA-seq data, we used unique detailed spatial maize kernel transcriptome (Zhan et al., 2015). Although different (sub)compartments and developmental stages (8 DAP vs 13 DAP) were used, we re-treated both RNA-seq raw data-set using the same bioinformatic pipeline and the same genome version (see Material and Methods) in order increase comparability. Hierarchical clustering showed that the transcriptomic signatures of our whole compartment samples (emb, End and Per) are close to the equivalent compartments generated by Zhan et al. (2015) *i.e* EMB (=embryo), CSE + CZ + AL (= three different endosperm sub-compartments), and PE (=pericarp) respectively (Supplemental Figure 2).

In summary, we have generated RNA-seq profiles from 13 DAP maize kernel compartments and embryo/endosperm interfaces. We have made this data available to the community in a user-friendly format via the eFP Browser (http://bar.utoronto.ca/efp_maize/cgi-bin/efpWeb.cgi?dataSource=Maize_Kernel) (*Available when publication will be accepted*).

### Preferentially expressed genes and biological processes associated with specific maize kernel (sub)compartments

Differential expression analyses were performed between the 6 (sub)compartments by comparing expression differences between pairs of tissues using a likelihood ratio test with *p*-values adjusted by the Benjamini-Hochberg procedure to control false discovery rates (see Material and Methods). Genes with both adjusted *p*-values lower than 0.05 and an expression difference of 4-fold or greater (log_2_(Fold Change) ≥ 2) were classed as differentially expressed genes (DEGs) (Supplemental Table 2). The full lists of DEGs for the 15 inter-tissue comparisons performed are available in Supplemental Data Set 2.

To identify the biological processes associated with the DEGs, a gene ontology (GO) analysis was performed. Due to the limited resources available, a new genome-wide annotation of all predicted proteins was carried out and linked to GO terms (see Material and Methods). In a first instance, GO terms enriched in the two zygotic compartments Emb and End were identified by analysing DEGs upregulated in each compartment compared to the two other main compartments (Table 1). The top ten GO terms enriched in the DEGs upregulated in the embryo relative to endosperm and pericarp, showed a significant enrichment in GO terms related to the cell cycle, DNA organization and cytoskeleton organization, consistent with the extensive developmental and mitotic activity within the embryo at this stage (Table 1). In contrast the GO terms enriched in the DEGs upregulated in the endosperm relative to embryo and pericarp were linked to metabolic functions such as nutrient reservoir activity, starch biosynthetic and metabolic processes (Table 1). These enrichments were consistent with the fact that the endosperm is a nutrient storage compartment where starch and reserve proteins are synthesized (Nelson and Pan, 1995; Zheng and Wang, 2015).

**Table 1:**
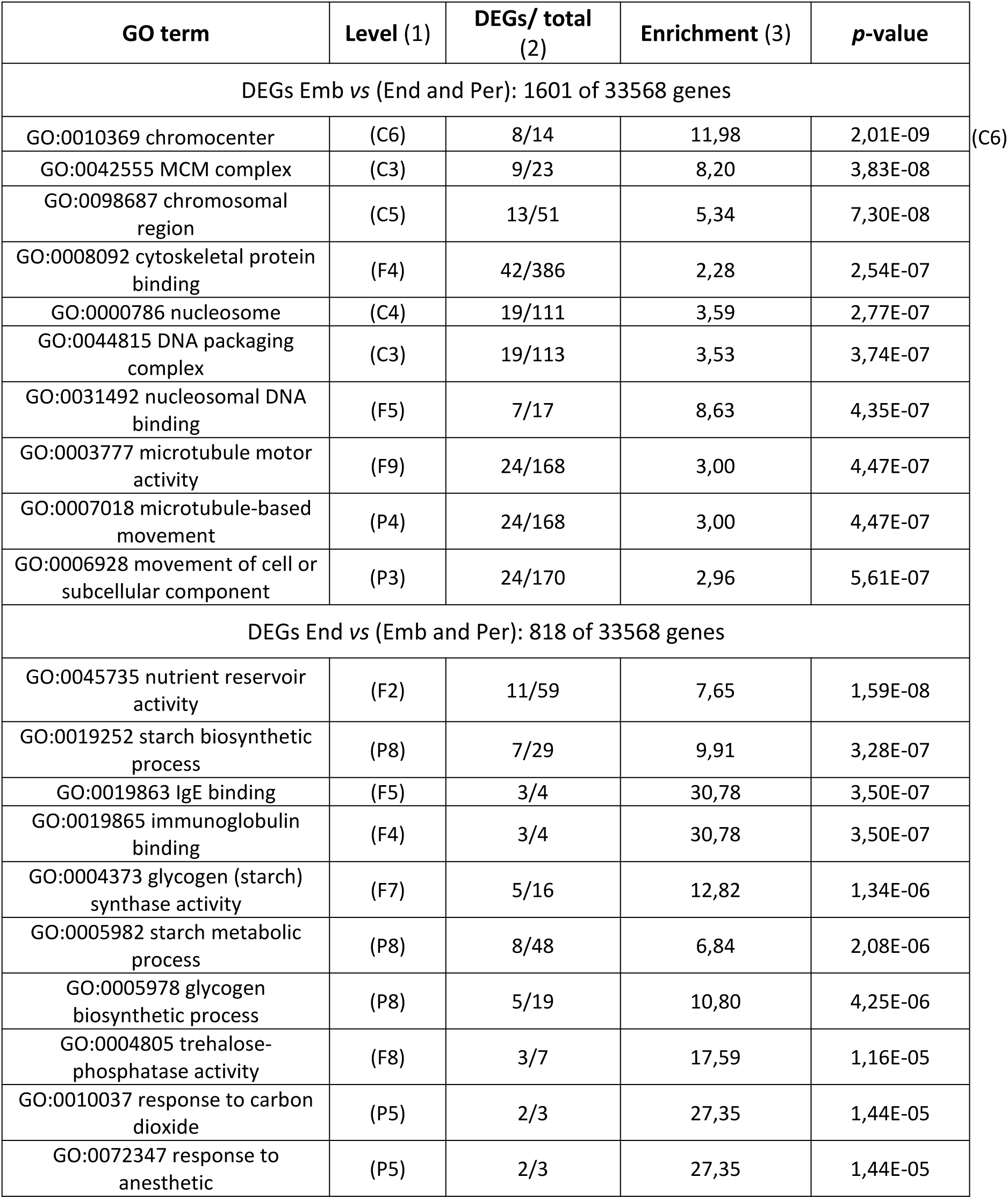
Top ten GO terms (sorted by increasing on p-value) enriched in the differentially expressed genes (DEGs) upregulated in one main compartment compared to the two others. Emb = embryo, End =endosperm, Per = pericarp (1) Minimal depth of the GO term in the GO tree, *‘P’* = biological process, ‘*F’*=molecular function and ‘*C’* = cellular component. (2) Number of genes associated with the GO term in the DEGs list / Number of GO term annotated genes in the maize genome. (3) The enrichment is defined in the Material and Methods.

### Enrichment for putative transporters at the endosperm/embryo interface

Focusing in on the embryo/endosperm interfaces, DEGs between the three sub-compartments (AS, SAL and EAS) and their whole compartments of origin were identified (Supplemental Table 2). 682 genes were found to be differentially expressed between AS and Emb according to the above criteria. Among them, 82 were more strongly and 600 more weakly expressed in AS compared to Emb samples (Supplemental Table 2). As expected, *ZmNAC124*, which is expressed in the coleorhiza (Figure 2B and Figure 3C,D) (Zimmermann and Werr, 2005) was found among the genes showing reduced expression in the apical scutellum. No GO terms were found to be significantly enriched in our analysis in the comparison of AS *vs* Emb (Table 2).

**Figure 3.**
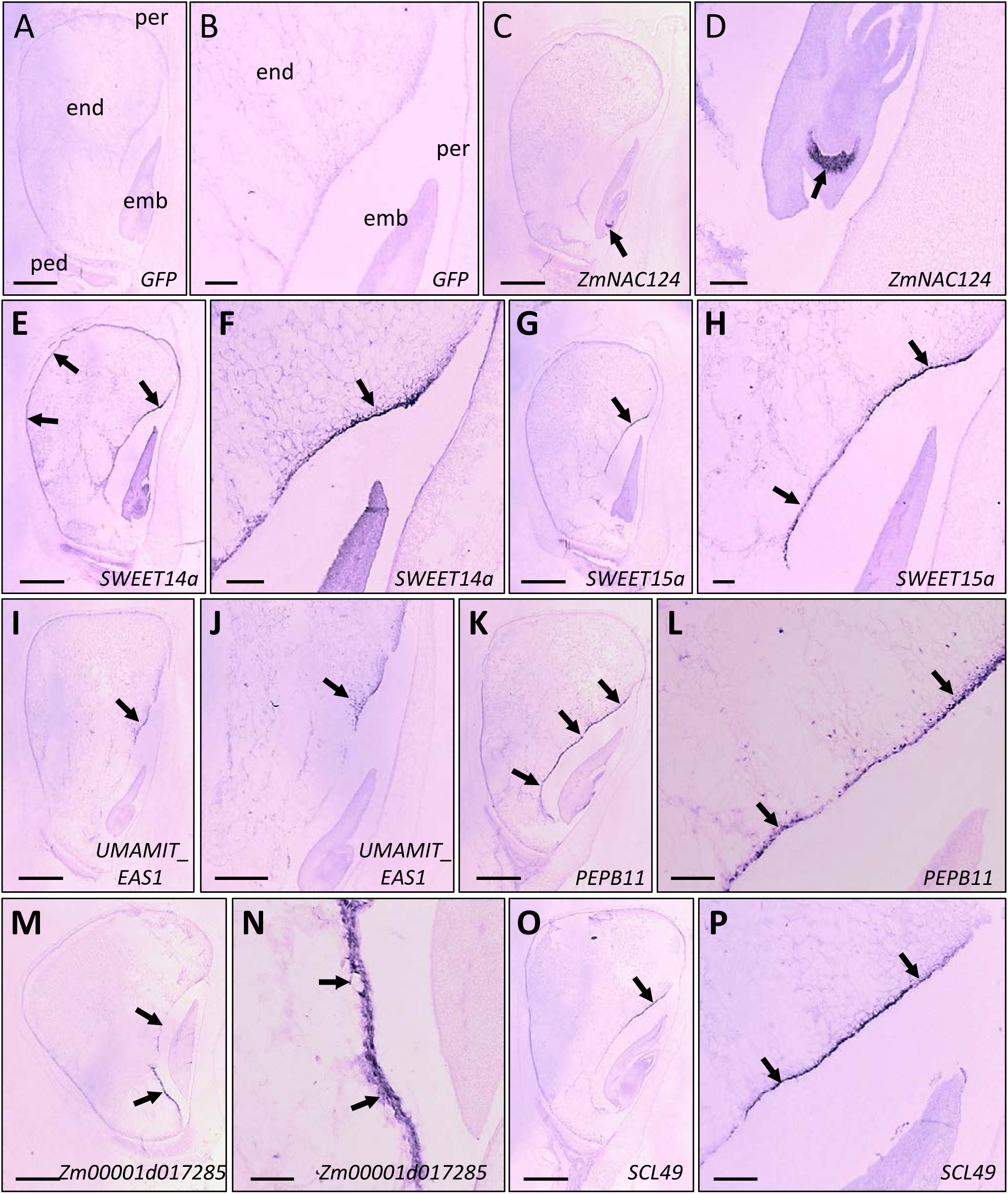
*In situ* hybridization on 13 DAP maize kernels probes detecting *GFP* (negative control) **(A, B)**, *ZmNAC124* (positive control) **(C, D)**, *SWEET14a* **(E, F)**, *SWEET15a* **(G, H)**, *UMAMIT_EAS1* **(I, J)**, *PEPB11* **(K, L)**, *Zm00001d017285* **(M, N)**, *SCL49* **(O, P)**. Scale bars correspond to 500 µm in A, C, E, G, I, J, K, M, O and 1000 µm in B, D, F, H, L, N, P. Arrows indicate main *in situ* hybridization signal. emb = embryo, end = endosperm, per = pericarp, ped = pedicel.

**Table 2:**
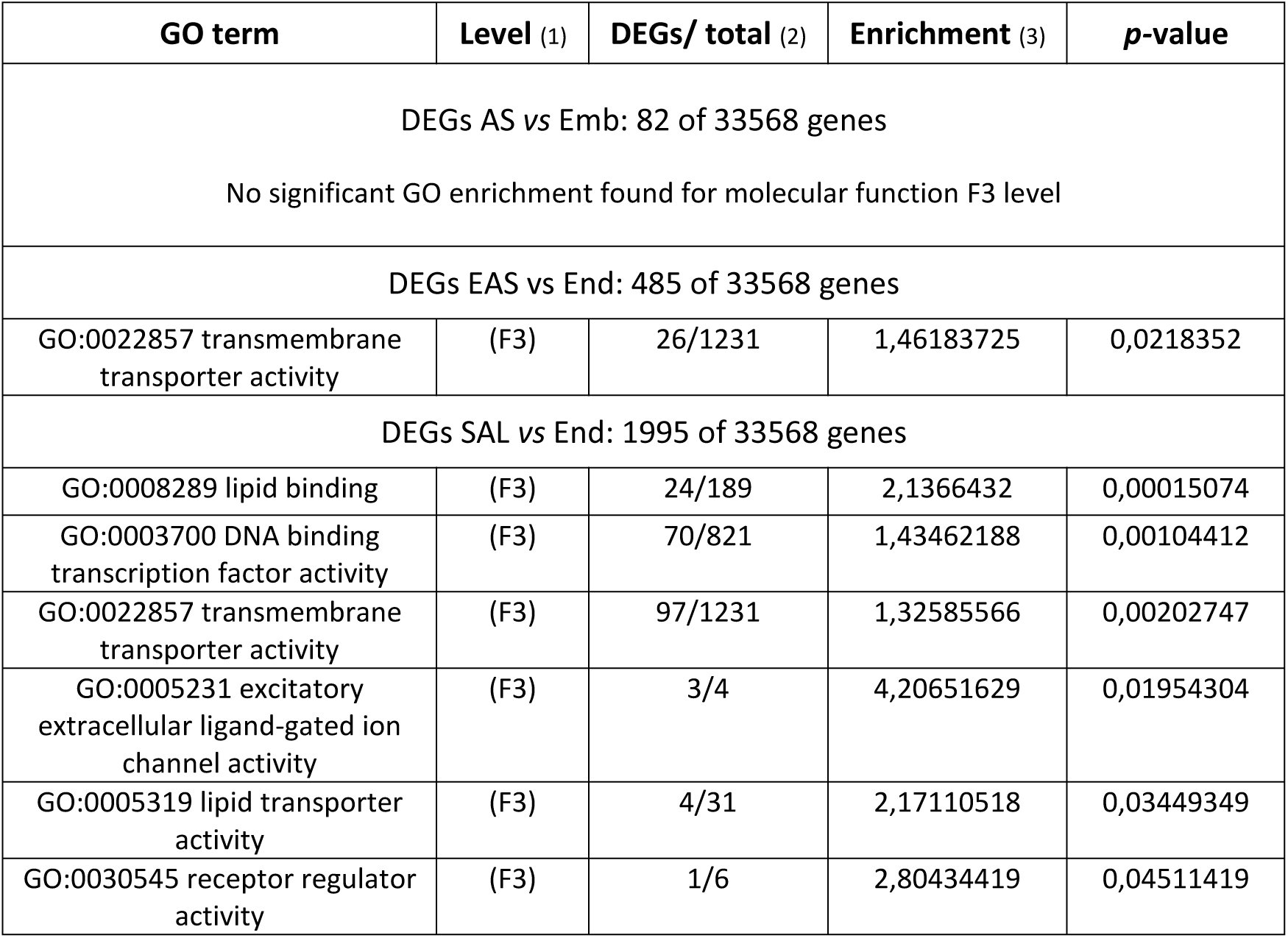
All GO terms from F3 (molecular function at level 3) significantly enriched in the differentially expressed genes (DEGs) upregulated in a sub-compartment compared to its compartment of origin. AS= Apical Scutellum, Emb = embryo, Embryo Adjacent to Scutellum (EAS), End =endosperm, and SAL = Scutellar Aleurone. (1) Minimal Depth of the GO term in the GO tree, *F stand for “*molecular function”. (2) Number of genes associated with the GO term in the DEGs list / Number of GO term annotated genes in the maize genome. (3) The enrichment is defined in the Material and Methods.

| GO term | Level (1) | DEGs/ total (2) | Enrichment (3) | p-value |
| --- | --- | --- | --- | --- |
| DEGs AS vs Emb: 82 of 33568 genes |  |  |  |  |
| No significant GO enrichment found for molecular function F3 level |  |  |  |  |
| DEGs EAS vs End: 485 of 33568 genes |  |  |  |  |
| GO:0022857 transmembrane transporter activity | (F3) | 26/1231 | 1,46183725 | 0,0218352 |
| DEGs SAL vs End: 1995 of 33568 genes |  |  |  |  |
| GO:0008289 lipid binding | (F3) | 24/189 | 2,1366432 | 0,00015074 |
| GO:0003700 DNA binding transcription factor activity | (F3) | 70/821 | 1,43462188 | 0,00104412 |
| GO:0022857 transmembrane transporter activity | (F3) | 97/1231 | 1,32585566 | 0,00202747 |
| GO:0005231 excitatory extracellular ligand-gated ion channel activity | (F3) | 3/4 | 4,20651629 | 0,01954304 |
| GO:0005319 lipid transporter activity | (F3) | 4/31 | 2,17110518 | 0,03449349 |
| GO:0030545 receptor regulator activity | (F3) | 1/6 | 2,80434419 | 0,04511419 |

The comparison between the EAS and the End revealed 1 498 DEGs with 485 genes showing stronger expression in the EAS than the End and 1 013 genes with the inverse profile (Supplemental Table 2). Among the genes more strongly expressed in the EAS, our GO analysis revealed a significant enrichment in only one GO term (GO analysis on molecular function terms at F3 level): “transmembrane transporter activity” (Table 2), which suggests a strong expression of transporter-encoding genes in the EAS.

Finally, 2 975 genes were found to be differentially expressed between SAL and End, 1 995 corresponding to genes more strongly expressed in the SAL, and 980 to genes with lower expression levels in the SAL (Supplemental Table 2). Interestingly, in the first group our GO analysis revealed an enrichment in three (out of 5) GO terms related to transport (Table 2).

A closer look at gene families encoding transporters amongst DEGs confirmed the overrepresentation seen in the GO analysis and revealed differences between the SAL and EAS. Among the genes that were at least 8 times more strongly expressed compared to End, 8.45 % (45/532) of the genes enriched in the SAL and 16.04 % (34/212) of the genes enriched in the EAS have at least one orthologue in rice or in Arabidopsis that encodes a putative transporter (Table 3). In the SAL, transcripts of genes encoding MATE (Multi-antimicrobial extrusion protein), which has been implicated in many diverse array of functions (for review see Upadhyay et al., 2019) and ABC (ATP-binding cassette) transporters were found to be the most strongly enriched, whereas in the EAS, genes encoding transporters from the MtN21/UMAMIT (Usually Multiple Acids Move In And Out Transporter), MtN3/SWEET (Sugars Will Eventually be Exported Transporter), and ABC transporter-families were the most represented. When looking at the putative molecules transported, a large number of genes encoding putative amino acid transporters was found to show stronger expression in the EAS than End samples, although genes encoding transporters for various other molecules including sugars, heavy metals, phosphate, inorganic ions or nucleotides also showed stronger expression (Table 3). Regarding the comparison of SAL vs End, mainly transporters putatively involved in amino acid and inorganic ions transport have been identified (Table 3). In summary, our work shows that both SAL and EAS cells strongly express putative transporter-encoding genes, suggesting that these cells are characterised by an elevated transmembrane transport of various molecules, and potentially mediate nutrient repartitioning around the embryo. However, each tissue preferentially expresses different classes of transporters, with MtN21/UMAMIT and MtN3/SWEET transporters involved respectively in amino acid or sugar transport more likely to be enriched in the EAS.

**Table 3:** Number of genes encoding putative transporters in the DEGs upregulated in the SAL or in the EAS compared to the End per family and per molecules putatively transported. Analysis was done base on orthology to rice and Arabidopsis (see material and method section).

| Transporter family | Ratio SAL/End > 8 | Ratio EAS/End > 8 |
| --- | --- | --- |
| MtN21/UMAMIT | 1 | 5 |
| MtN3/SWEET | 0 | 3 |
| AAP | 1 | 2 |
| MATE | 7 | 1 |
| ABC | 3 | 4 |
| GDU | 1 | 2 |
| VIT | 0 | 2 |
| Phosphate transporters | 0 | 2 |
| Other | 32 | 13 |
| Total number | 45 | 34 |
| % in the gene list | 8.45% | 16.04 % |
| **Molecules putatively transported** | **Ratio SAL/End > 8** | **Ratio EAS/End > 8** |
| Amino acids and/or auxin | 7 | 12 |
| Nucleotides | 1 | 1 |
| Heavy metal | 3 | 3 |
| Sugar | 0 | 4 |
| Phosphate | 0 | 2 |
| Other inorganic ions | 5 | 2 |

### The EAS is restricted to one to three endosperm cell layers adjacent to the scutellum

The SAL has both cellular and biochemical characteristics of the aleurone making it inherently different from other endosperm tissues (Gontarek and Becraft, 2017; Zheng and Wang, 2014). In contrast, EAS cells have not been reported to have distinct features that would allow to distinguish them cytologically from SE cells, which compose the majority of the volume of the endosperm (Van Lammeren, 1987). However, our transcriptomic analysis suggests that these cells deploy a specific genetic program. In order to (1) confirm EAS expression specificity and to (2) provide a more precise spatial resolution to define and characterize this new region, *in situ* hybridizations were performed with a set of 6 genes more than 10 fold enriched in the EAS transcriptome compared to the End transcriptome (Supplemental Table 3, and Supplemental Figure 3 for two examples of eFP browser pattern). Three of these genes encode putative transporters, namely *SWEET14a* (*Zm00001d007365*) and *SWEET15a* (*Zm00001d050577*) encoding putative sugar transporters of the SWEET family, and *Zm00001d009063* called *UMAMIT*_*EAS1* encoding a putative amino acid transporter belonging to the *UMAMIT* family (Müller et al., 2015; Sosso et al., 2015). The three remaining genes were *PHOSPHATIDYLETHANOLAMINE-BINDING PROTEIN 11* (*PEBP11*, *Zm00001d037439*), *SERINE CARBOXYPEPTIDASE-LIKE 49* (*SCL49*, *Zm00001d014983*) and *Zm00001d017285,* a gene with no name and unknown function (Supplemental Table 3). The negative control chosen for *in situ* hybridizations, was an antisense probe generated against a GFP-encoding ORF. The positive control was *ZmNAC124* (*Zm00001d046126*), which is specifically expressed in the Emb compartment in our transcriptome (Figure 2B and Supplemental Table 3) and which had previously been shown by *in situ* hybridization to be expressed in specific embryonic tissues (Zimmermann and Werr, 2005). *In situ* hybridizations were performed on 13 DAP kernels, the same stage as used for the transcriptome analysis. The 4 probes detecting *SWEET15a* (Figure 3G, H), *PEPB11* (Figure 3K, L), *Zm00001d017285* (Figure 3M, N) and *SCL49* (Figure 3O, P) gave a strong signal restricted to a few layers of endosperm cells immediately adjacent to the scutellum, with little or no expression detected elsewhere in the kernel. For the probe directed against *SWEET14a* the signal was strong in the EAS, but was also present, albeit more weakly, in other kernel tissues, especially in the embryo and aleurone (Figure 3E, F). The probe against *UMAMIT-EAS1* gave a weaker signal restricted to the apical part of the EAS region, consistent with the lower expression levels of this gene in our transcriptome data (Supplemental Table 3). However, the signal for *UMAMIT-EAS1* was specific to these EAS cells (Figure 3 I, J). These results confirmed that EAS cells have a specific transcriptional program and that this programme (and thus the EAS) is restricted to 1 to 3 layers of endosperm cells adjacent to the scutellum as confirmed by sagittal sections (Figure 5A-B).

### The EAS is a dynamic region reflecting the period of strong embryo growth

To evaluate the dynamics of gene expression in the EAS during kernel development, *in situ* hybridizations were carried out on kernels at different developmental stages (9, 11, 14, 17 and 20 DAP) (Figure 4 and Supplemental Figure 4). The four probes giving a strong and EAS-specific signal at 13 DAP (*SWEET15a, PEPB11, Zm00001d017285, SCL49*) were used (Figure 4). In 9 DAP kernels, the probes for *PEPB11* and *SCL49* showed no signal, whereas those for *SWEET15a* and *Zm00001d017285* gave a strong signal in the endosperm cells adjacent to the apical part of the embryo (Figure 4 and Supplemental Figure 4). This signal was restricted to the few layers of cells in the vicinity of the nascent scutellum. At this stage, the basal part of the embryo was still surrounded by ESR cells and no signal was detected in this region. At 11 DAP, all four probes tested gave a very strong signal in the layers of endosperm cells adjacent to the scutellum. At 14 DAP and 17 DAP, the signal was still detected and restricted to the cell layers in close contact with the embryo (Figure 4 and Supplemental Figure 4). Finally, at 20 DAP the signal decreased for all four probes with a total disappearance for *SWEET15a*. Together, these results revealed that the EAS transcriptomic region was restricted to a defined time window. Its onset at 9 DAP was concomitant with the formation of the scutellum, marking a switch in embryo/endosperm interactions from an ESR/embryo to an EAS/scutellum interface. Its decline occurred around 20 DAP when rapid embryo growth comes to an end.

**Figure 4.**
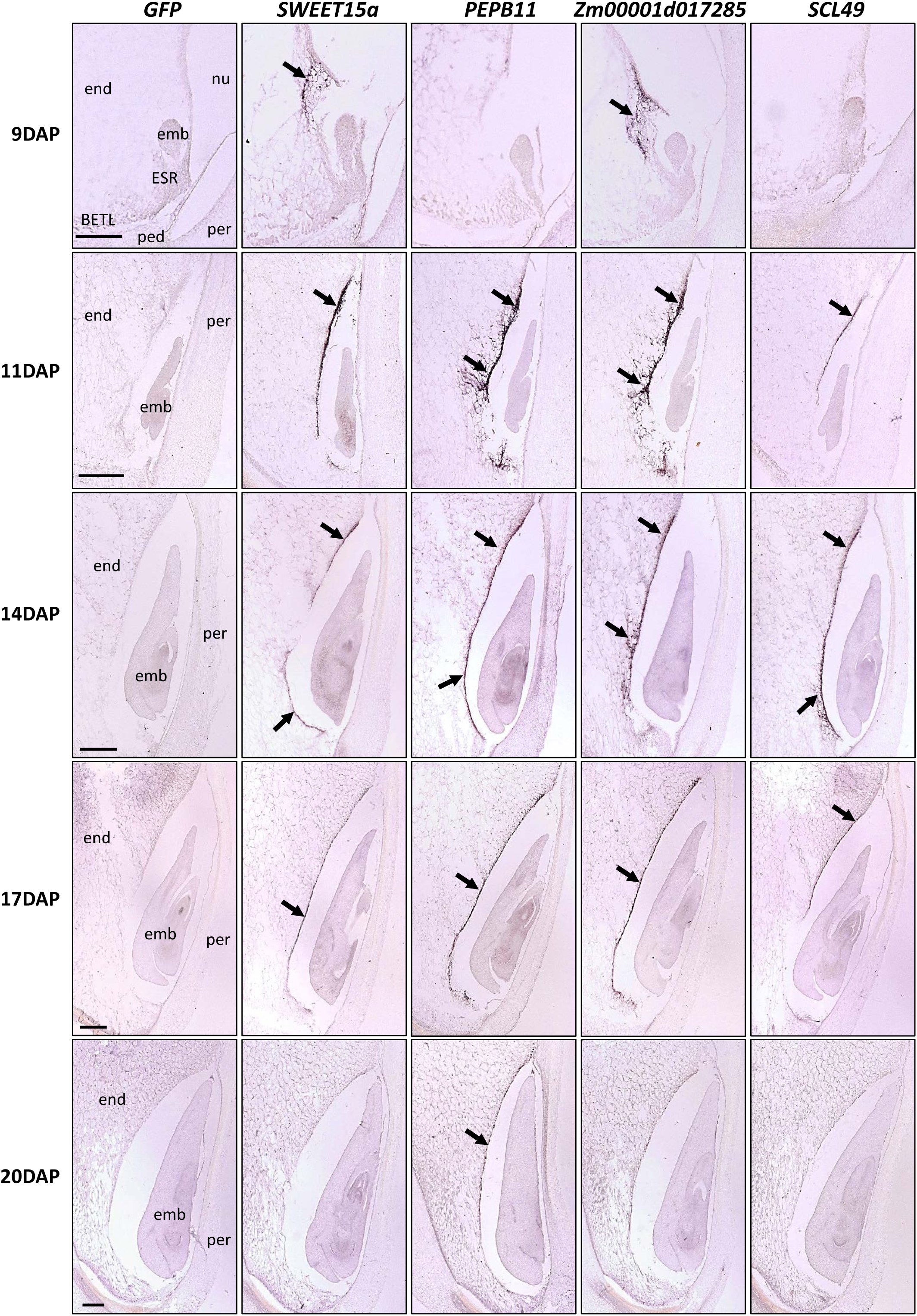
*In situ* hybridization of 4 probes detecting EAS marker genes (*SWEET15a*, *PEPB11*, *Zm00001d017285*, *SCL49*) on kernel sections at different developmental stages. Pictures are zoom from Supplemental Figure 4, and scale bars correspond to 200 µm for 9 DAP kernels and 500 µm for the other stages. For each image the name of the probe is indicated at the top of the figure and the stage on the left. Arrows indicate main *in situ* hybridization signal. End = endosperm, emb = embryo, per = pericarp, nu = nucellus, ESR = embryo surrounding region, BETL = basal endosperm transfer layer, ped = pedicel.

### EAS cells originate from the starchy endosperm (SE) and undergo cell death

Despite the preferential or specific expression of EAS marker genes, and consistent with their SE-like morphology, EAS cells also showed some transcriptomic characteristics of the SE such as a strong expression of genes encoding zein storage proteins (Supplemental Figure 5). The presence of *ZEIN* transcripts in the EAS region is supported by *in situ* hybridization data (Woo et al., 2001). In order to perform a more global comparison, we asked to which samples from the Zhan et al (2015) dataset our EAS transcriptome was most similar (Supplemental Figure 2). EAS was more similar to the two SE sub-regions: central starchy endosperm (CSE) and conducting zone (CZ) than to other samples, strengthening the idea that EAS originate from the starchy endosperm.

To address the question of EAS cell fate in proximity to the scutellum, sagittal sections of the EAS/scutellum interface were both hybridized with a EAS-specific probes (against *SWEET15a*) and stained with calcofluor to reveal cell walls (Figure 5A-B). Accumulation of cell wall material occurred at the endosperm interface with the scutellum, which may result from the compaction of crushed endosperm cells. Interestingly, *in situ* hybridization signal for the EAS marker genes was found in the first uncrushed cell layer (Figure 5B). An appealing model is that EAS cells are actually SE cells that are forced into juxtaposition with the scutellum because of the invasive growth of the embryo into the SE during kernel development (Figure 6), suggesting that the EAS program may not be fixed within a static group of cells but instead be triggered as SE cells enter into contact with the scutellum.

**Figure 5.**
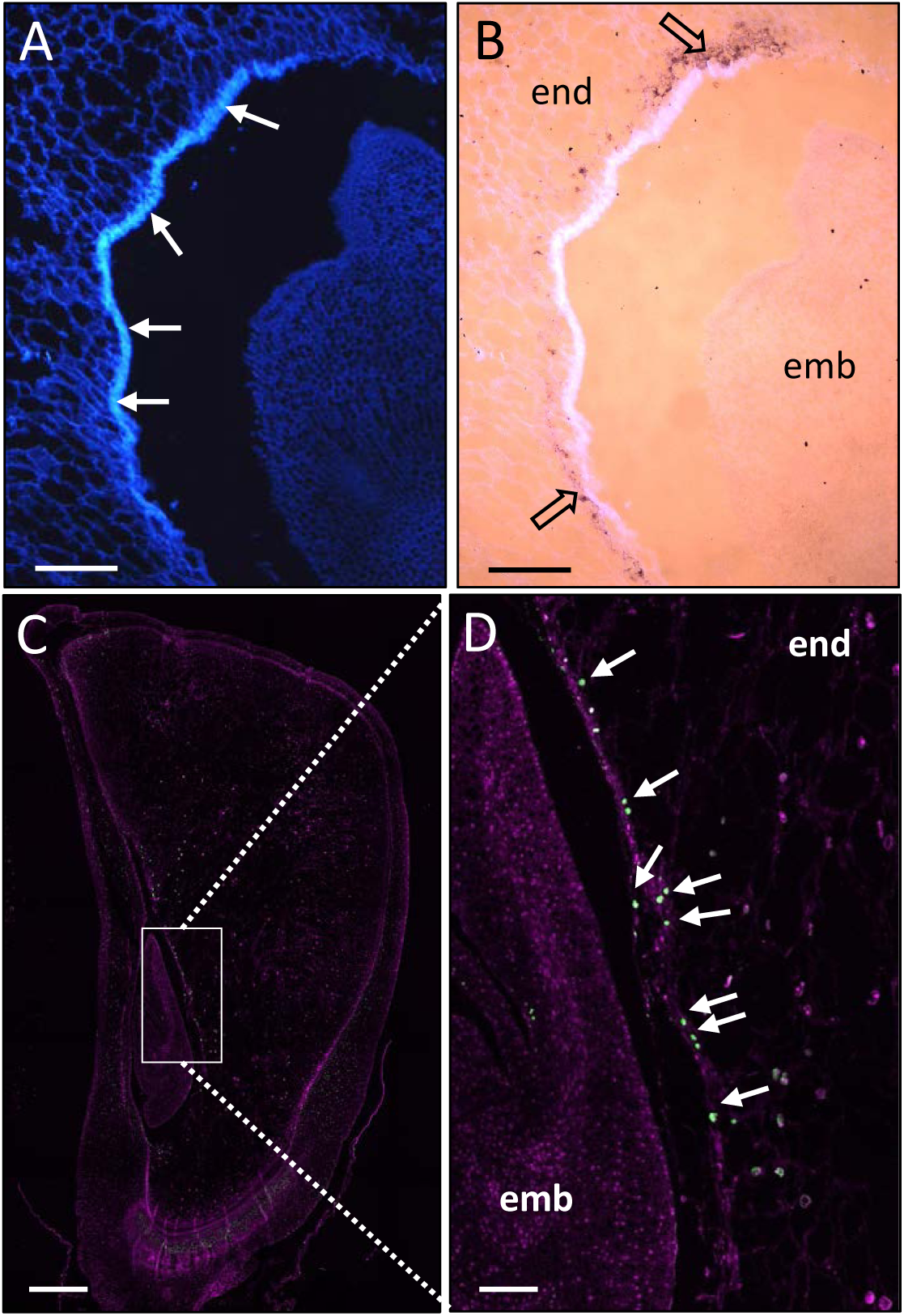
Crushed cell walls and cell death occurs in the EAS. **(A,B)** Calcofluor staining of cell walls of 13 DAP maize kernel sections **(A)** together with *in situ* hybridization with *SWEET15a* antisense probes **(B)** on sagittal section. Empty black arrow indicates *in situ* hybridization signal, while plain white arrows indicate the accumulation of crushed cell walls. **(C,D)** TUNEL labelling of 15 DAP kernels. Fluorescein labelling of the TUNEL positive nuclei is shown in green and propidium iodide counterstaining in purple. Arrows indicate the nucleus stained by TUNEL in the EAS. Scale bars correspond to 200 µm in (A,B) and 500 µm in (C) and 100 µm in (D). emb = embryo, end = endosperm.

**Figure 6.**
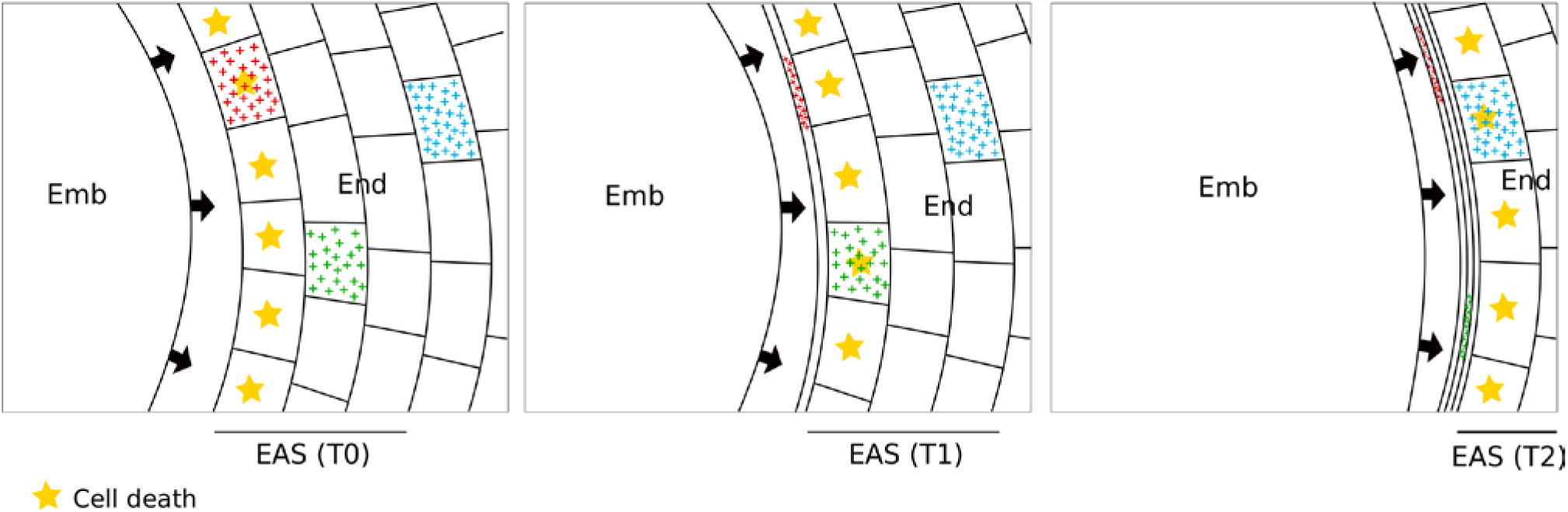
Scheme summarizing the EAS dynamic. Three different consecutive times points (t0, t1 and t2) are represented. Embryo scutellum invades (representing by arrows) the surrounding endosperm which enters in cell death (yellow stars). Three cells are labelled by a cross pattern to illustrates this dynamic. Emb = embryo scutellum, End = endosperm, EAS = endosperm adjacent to scutellum.

If this model is correct, EAS cells would be likely to be successively eliminated as they come into contact with the embryo. Terminal deoxynucleotidyl transferase dUTP Nick End Labelling (TUNEL) assays were performed on 14 DAP kernels to visualize DNA degradation as an potential indicator of the presence of dying cells (Labat-Moleur et al., 1998). In addition to the PC and coleoptile regions both previously reported to give strong TUNEL signals (Giuliani et al., 2002; Kladnik et al., 2004), a clear TUNEL positive signal was also found in some EAS cells in close contact with the scutellum (Figure 5C, D). This result is consistent with the possibility that a form of cell death occurs at this interface. To clarify whether transcriptional activation of EAS specific genes precedes or succeeds cell death, the expression levels of genes associated with programmed cell death in plants were analysed (Supplemental Figure 6) (Arora et al., 2017; Fagundes et al., 2015). Surprisingly none of the previously identified programmed cell-death associated genes was found to be particularly up regulated in the EAS compared to End or other samples. In addition, no enrichment of GO terms associated with programmed cell death was found in the DEGs strongly expressed in the EAS relative to the End samples (Table 2). These data suggested that either only a small proportion of EAS cells undergo cell death or crushing of EAS cells does not trigger “classical” programmed cell death programmes. A parallel could be made with accidental cell death (ACD) defined in animal, in which cells are dying as a result of their immediate structural breakdown due to physicochemical, physical or mechanical cues (Galluzzi et al., 2015).

### Impaired expression of some EAS marker genes in *emb* mutants

To test to what extent the proximity of the embryo/scutellum was required for EAS gene expression, the *embryo specific* mutation *emb8522* was used in an Rcsm2 genetic background enhancing the early embryo deficient phenotype (Sosso et al., 2012). In this background the recessive *emb8522* mutation produced vestigial embryos composed of a small heap of cells. Nevertheless, a cavity corresponding to the size a normal embryo was generated that was only very partially occupied by the aborting embryo (Heckel et al., 1999; Sosso et al., 2012). Self-fertilization of heterozygous plants carrying the *emb8522* mutation was performed, and *in situ* hybridizations were carried out on 13 DAP sibling kernels with either phenotypically wild-type or mutant embryos, to visualize the transcripts of four EAS marker genes (Figure 7). Similar EAS specific expression patterns were observed in Rcsm2 kernels with embryos (Figure 7) to those observed in B73 kernels (Figure 3 and 4) for all genes tested, indicating a conservation of EAS cell identity in this genetic background. In *emb* kernels, the probes detecting *Zm00001d017285* and *SWEET15a* still showed a signal in the EAS region but with an altered distribution (Figure 7). In *emb* kernels, *Zm00001d017285* expression was found to be restricted to the apical part of the embryo cavity and *SWEET15a* expression expanded to the SAL, suggesting an inhibitory role of the normal embryo on *SWEET15a* expression in this tissue. Interestingly the two other EAS marker genes tested showed either only very weak expression (*SCL49*) or no expression (*PEPB11*) in *emb* kernels, indicating a promoting effect of the normal embryo on the expression of these two genes (Figure 7).

**Figure 7.**
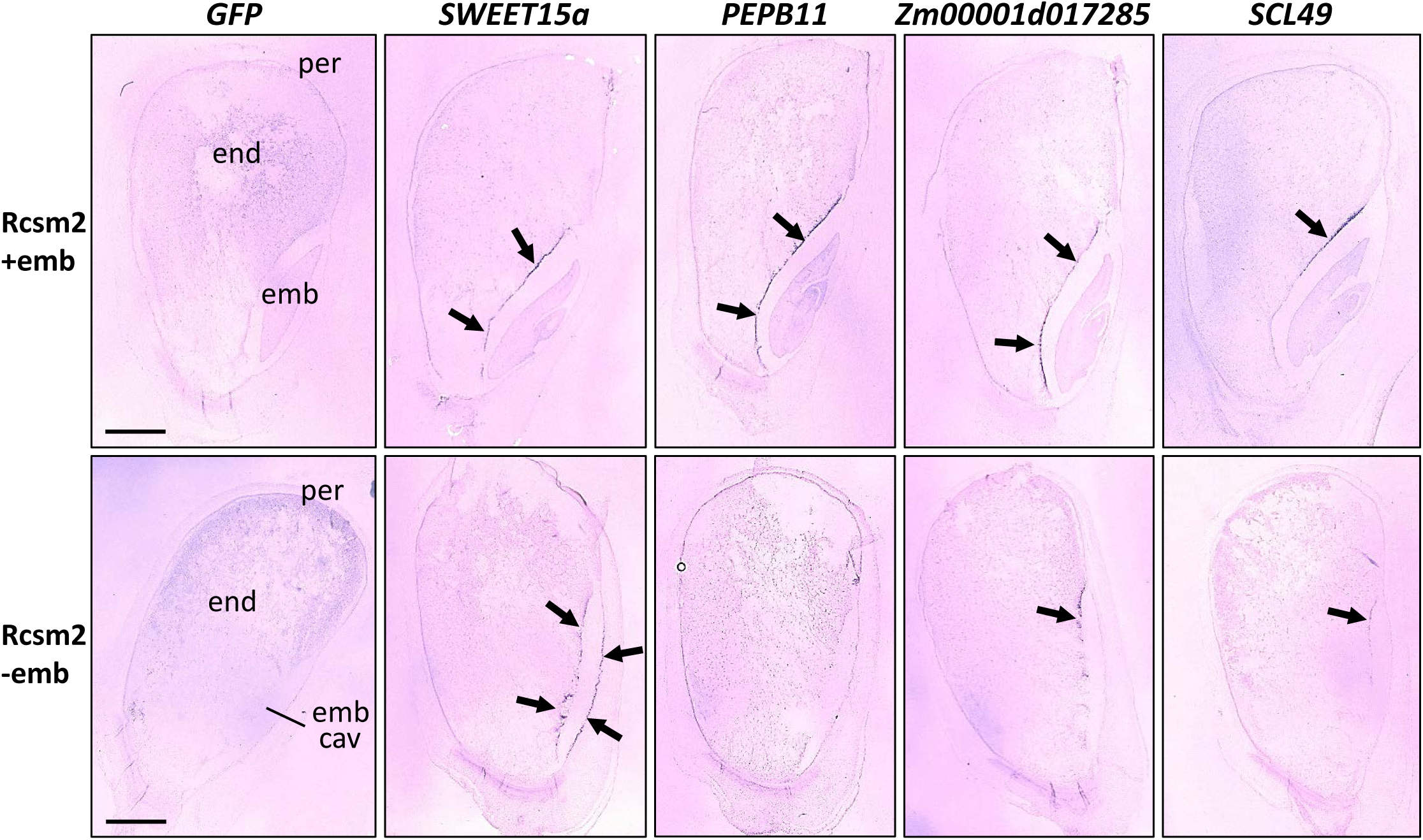
*In situ* hybridization with several probes marking the EAS on 13 DAP maize kernel sections of the Rcsm2 genetic background. Kernels come from a self-pollination of a mother plant heterozygous for the *emb8522* mutation. Upper line corresponds to kernels with embryo (*emb8522* +/- or +/+), and the bottom line to kernels without embryo (*emb8522 -/-)*. Arrows indicate the main *in situ* hybridization signal. Scale bars correspond to 1000 µm.

## Discussion

### Transcriptomes at embryo/endosperm interfaces

As in other flowering plants, seed development in maize is governed by specific temporal and spatial genetic programs, distinguishing early development, filling and maturation on one hand and embryo, endosperm and pericarp on the other (Chen et al., 2014; Downs et al., 2013; Li et al., 2014; Lu et al., 2013; Meng et al., 2018; Qu et al., 2016). Recently a transcriptome analysis on nucellus (including the fertilized embryo sac) increased the temporal resolution and allowed unprecedented access to information regarding the genetic control of early seed developmental (Yi et al., 2019). The most detailed spatial analysis to date used Laser-Capture Microdissection (LCM) on 8 DAP kernels (Zhan et al., 2015) to reveal the expression of specific populations of genes in the maternal tissues, the embryo and the main endosperm cell types namely ESR, BETL, AL and SE (which was subdivided in CSE and CZ). Although providing an extremely valuable resource, this study did not address the question of whether specific transcriptional domains exist at the embryo-endosperm interface.

Both the endosperm and embryo are complex compartments with several morphologically and functionally distinct domains (Olsen, 2004b; Sabelli and Larkins, 2009). Because they undergo complex and coordinated developmental programmes, the interfaces between the embryo and the endosperm represent important, and constantly changing zones of exchange, both in term of nutrition and communication (Ingram and Gutierrez-Marcos, 2015; Nowack et al., 2010; Widiez et al., 2017). In order to understand these interactions two subdomains of the endosperm and one subdomain of the embryo were hand-dissected: the scutellar aleurone layer (SAL) at the adaxial side of the embryo, a band of several cell layers of the starchy endosperm at the abaxial side of the embryo and the scutellum of the embryo. 13 DAP kernels were chosen for our analysis because at this stage the embryo has emerged from the ESR and is establishing new interactions with endosperm. From a practical point of view, 13 DAP is also the earliest stage allowing reliable hand dissection of the chosen interfaces.

Contamination with neighbouring tissues is an important issue in any dissection experiment. For example, in Arabidopsis, an extremely valuable and globally very reliable resource generated by LCM (Belmonte et al., 2013; Le et al., 2010) was recently shown to contain some inter-compartment contamination, which caused problems for the investigation of parental contributions to the transcriptomes of early embryos and endosperms (Schon and Nodine, 2017). In our study, precautions were taken to limit inter-tissue contamination by (i) washing each sample before RNA extraction (see material and methods) and (ii) generating four biological replicates for each tissue. Marker gene analysis confirmed the conformity of the samples with the exception of a potential minor contamination of pericarp by the AL, suggested by the apparent expression of both the *AL9* and Zm00001d024120 aleurone marker genes and the endosperm marker genes *ZmZOU/O11* and *O2* in the pericarp sample (Figure 2C, D). This could have been caused by the tendency of the AL to stick either to the starchy endosperm or to the pericarp.

### The EAS, a novel endosperm subdomain likely involved in carbon and nitrogen fluxes from the endosperm to the embryo

Transcriptomic profiling of the two endosperm interfaces with the embryo (SAL and EAS) revealed specific transcriptional signatures. While this could have been expected for the cytologically distinct SAL, it was rather unexpected for the cell layers adjacent to the abaxial side of the embryo, which do not present any obvious cytological differences to other SE cells (Van Lammeren, 1987). Based on the observed enrichment of hundreds of transcripts in these cell layers, they represent a novel subdomain of the endosperm which we named “endosperm adjacent to scutellum” (EAS).

GO analysis revealed a significant enrichment in the GO category transmembrane transporter activity for both the SAL and EAS, and additionally for lipid transporter activity and ion channel activity for SAL (Table 2). A closer look at DEGs for both EAS and SAL shows the presence of different transporter gene families (Table 3). Interestingly, many *UMAMITs* and *SWEETs*, thought to transport amino acids/auxin and sugars, respectively, were found enriched in the EAS. UMAMITs and SWEETs are considered to be bi-directional transporters, although they tend to act as exporters when located at the plasma membrane, exporting nutrients down concentration gradients generated by sinks in adjacent tissues (Chen et al., 2012; Müller et al., 2015). Two non-exclusive hypotheses could explain the elevated expression of transporter-encoding genes in the EAS: either these cells actively take up nutrients that arrive from the BETL via the SE and then export them into the apoplastic space surrounding the growing embryo, or they are simply involved in recycling nutrients from dying endosperm cells that are crushed by the growing embryo.

With regard to nutrient uptake on the embryo side, one might expect the expression of genes encoding nutrient importers at the surface of the scutellum in order to take up apoplastic metabolites. However, in our apical scutellum transcriptome (AS) we were not able to detect differentially expressed importer-encoding genes with respect to the entire embryo (Emb). While this could suggest that the regulation of importer activity does not occur at the transcriptional level, it seems more likely that our transcriptomic comparison AS *vs* Emb was not well designed for the identification of such genes, since the whole embryo is mainly composed of scutellum tissues.

In the future a more detailed comparison of the gene expression profiles of the BETL (import) and the EAS (export) regions could be informative. The BETL is an interface specialized in nutrient transfer from maternal phloem terminals to the endosperm (Chourey and Hueros, 2017). The hexose transporter SWEET4c is preferentially expressed in the BETL, and loss of function of *SWEET4c* results in the production of a shrivelled endosperm, illustrating the critical importance of hexose transport in the BETL for normal endosperm growth (Sosso et al., 2015). Interestingly, *SWEET4c* is also found in the DEGs showing strong expression in the EAS compared to the endosperm as a whole, possibly suggesting commonalities between BETL and EAS function. EAS-specific knock-down of *SWEET4c* might be one strategy to test this hypothesis and to address the question of possible redundancy with *SWEET14a* and *SWEET15a,* also enriched in the EAS. Nonetheless, notable differences exist between the EAS and the BETL. Firstly BETL cells have structural features including dramatic cell wall ingrowths that make them unique in the endosperm (Chourey and Hueros, 2017; Leroux et al., 2014). In contrast EAS cells cannot be morphologically differentiated from the SE (Van Lammeren, 1987). Secondly, the BETL represents a static interface, contrary to the EAS which is displaced as the embryo scutellum expands (Figure 6) during to the most rapid growth-phase of the embryo (Chen et al., 2014).

### The EAS is a developmentally dynamic interface

The detection of DNA fragmentation, a characteristic of cell death, in EAS cells (Figure 5C, D), together with an accumulation of cell wall material in this zone (Figure 5A, B) suggested that endosperm cells are eliminated as the embryo grows. An important question is whether this involves a genetically controlled cell-autonomous death or a more atypical and passive cell death process caused by embryo growth. In the Arabidopsis seed, where most of the endosperm degenerates during seed development, the expression of developmental cell death marker genes such as *PLANT ASPARTIC PROTEASE A3 (PASPA3)* or *BIFUNCTIONAL NUCLEASE 1 (BFN1)* has been detected at the embryo interface (Fourquin et al., 2016; Olvera-Carrillo et al., 2015). In maize, where less is known about molecular actors involved in developmental cell death, a survey of putative cell death marker genes showed expression of most of these genes in EAS cells but no enrichment compared to the End (Supplemental Figure 6 and Supplemental Data Set 2). Although cell death in the EAS could be triggered by the activation of unknown cell death-associated genes, a more likely explanation for our observations could be a dilution of the transcriptional signal in the EAS transcriptome, making it undetectable. This is supported by TUNEL staining, which revealed a very localised signal limited to a few cells at the immediate interface with the embryo (Figure 5C, D). In addition, previous cell death staining with Evans blue, did not reveal any massive cell death in the EAS, further supporting the hypothesis of very localized cell death events (Young and Gallie, 2000).

The precise spatial organisation of cell death and transporter expression remains unclear but the expression of transporters might allow the recycling of nutrients from the cells before they die. As these cells are SE in origin, they could already have initiated nutrient storage at 13 DAP, as illustrated by substantial expression of *Zein* genes (Supplemental Figure 5). Nutrient recycling could be an advantageous way for the plant to efficiently reuse stored nutrients. Interestingly, in Arabidopsis the STP13 sugar transporter is upregulated in several cell death contexts but the role of transporters in nutrient recycling remains poorly documented in plants (Norholm et al., 2006).

### The importance of the embryo for the expression of EAS marker genes

Since the EAS is a mobile interface, forming adjacent to the expanding scutellum, we asked whether the presence/absence of the embryo influences the activation of EAS marker genes (Figure 7). In *8522emb* mutant kernels which produce a seemingly empty, but normally sized embryo cavity containing an aborted embryo (Heckel et al., 1999; Sosso et al., 2012) the expression of different EAS marker genes was affected in different ways. The *SWEET15A* gene was still expressed in EAS cells, but also became strongly expressed at the opposite embryo/endosperm interface (SAL). Based on the precedent of the SWEET4c transporter, which is induced by sugar (Sosso et al., 2015), it is possible that a similar induction could occur in the case of *SWEET15a*. The absence of a normal embryo could lead to a build-up of sucrose in the embryo cavity of *8522emb* mutants leading to such an induction. In contrast, the expression domain of the *Zm00001d017285* marker gene is reduced in *8522emb* mutants, with expression becoming restricted to the apical part of the EAS. Finally, the expression of *PEPB11* and *SCL49* is dramatically reduced in *8522emb* mutants compared to phenotypically wild-type kernels. Our results suggest that EAS-specific gene expression could be a result of several independent factors, some of which could originate from the endosperm, and others from the embryo. The mechanisms involved in embryo cavity formation remain elusive, although a recent study showed that the SHOHAI1 protein is required in the endosperm for the formation of the embryo cavity (Mimura et al., 2018).

Interestingly, the expression of both *PEPB11* and *SCL49* initiates relatively late in the EAS, whereas the expression of *SWEET15a and Zm00001d017285* initiates before 9 DAP. This suggests the presence of at least two transcriptional programs in the EAS: one initiating early and weakly influenced by the embryo and a second activated later, and more strongly embryo-dependent. The generation of comparable transcriptomes at earlier developmental stages could help us identify the key signals activating gene expression in the EAS, and potentially pinpoint transcription factors regulating gene expression in this tissue. In parallel, phenotypic analysis of loss-of-function mutants of genes enriched in the EAS is needed to further elucidate the biological role of this novel endosperm subdomain.

## Material and Methods

### Plant material, plant growth conditions

A188 and B73 inbred lines were cultivated in the green house as described previously (Rousseau et al., 2015). The *emb8522* mutant in the *rcsm-2* background (Sosso et al., 2012) and the B73 plants used for *in situ* hybridization were grown in a field plot located at the ENS de Lyon, France.

### Isolation of maize kernel compartments

Kernel (sub)compartments of the B73 inbred line were hand dissected and quickly washed with HyClone Dulbecco’s phosphate-buffered saline solution (ref. SH30378.02), before freezing them in liquid nitrogen. For each (sub)compartment, four independent biological replicates were produced (Supplemental Table 1). For each biological replicate, the material comes from two independent, 13 day-old maize ears, i. e. a total of 8 different ears was used for each (sub)compartment. Within each biological replicate, tissues from 4 to 84 kernels were pooled depending on the size of the considered (sub)compartment (Supplemental Table 1).

### RNA extraction and RNA-seq

Total RNAs were extracted with TRIzol reagent, treated with DNase using the Qiagen “RNase-Free DNase Set”, and purified using Qiagen RNeasy columns according to the supplier’s instructions. RNA-seq libraries were constructed according to the “TruSeq_RNA_SamplePrep_v2_Guide_15026495_C” protocol (Illumina®, California, USA). Sequencing was carried out with an Illumina Hiseq2000 at the IG-CNS (Institut de Génomique-Centre National de Séquençage). The RNA-seq samples were sequenced in paired-end (PE) mode with a sizing of 260 bp and a read length of 2×100 bases. Six samples were pooled on each lane of a HiSeq2000 (Illumina), tagged with individual bar-coded adapters, giving approximately 50 million paired-end (PE) reads per sample. All steps of the experiment, from growth conditions to bioinformatics analyses, were managed in the CATdb database (Gagnot et al., 2008, http://tools.ips2.u-psud.fr/CATdb/) with project ID “NGS2014_21_SeedCom” according to the MINSEQE (minimum information about a high-throughput sequencing experiment) standard (http://fged.org/projects/minseqe/).

### RNA-seq read processing and gene expression analysis

RNA-seq reads from all samples were processed using the same pipeline from trimming to counts of transcripts abundance as follows. Read quality control was performed using the FastQC (S. Andrew, http://www.bioinformatics.babraham.ac.uk/projects/fastqc/). The raw data (fastq files) were trimmed using FASTX Toolkit version 0.0.13 (http://hannonlab.cshl.edu/fastx_toolkit/) for Phred Quality Score > 20, read length > 30 bases, and ribosomal sequences were removed with the sortMeRNA tool (Kopylova et al., 2012).

The genomic mapper TopHat2 (Langmead and Salzberg, 2012) was used to align read pairs against the *Zea mays* B73 genome sequence (AGP v4, (Jiao et al., 2017)), using the gene annotation version 4.32 provided as a GFF file (Wang et al., 2016). The abundance of each isoform was calculated with the tool HTSeq-count (Anders et al., 2015) that counts only paired-end reads for which paired-end reads map unambiguously one gene, thus removing multi-hits (default option union). The genome sequence and annotation file used was retrieved from the Gramene database (http://www.gramene.org/ release 51, in September 2016 (Gupta et al., 2016)).

Choices for the differential analysis were made based on Rigaill et al., 2018. To increase the detection power by limiting the number of statistical tests (Bourgon et al., 2010) we performed an independent filtering by discarding genes which did not have at least 1 read after a count per million analysis (CPM) in at least one half of the samples. Library size was normalized using the method trimmed mean of M-values (TMM) and count distribution was modelled with a negative binomial generalized linear. Dispersion was estimated by the edgeR package (version 1.12.0, McCarthy et al., 2012) in the statistical software ‘R’ (version 2.15.0, R Development Core Team, 2005). Pairwise expression differences were performed using likelihood ratio test and *p*-values were adjusted using the Benjamini-Hochberg (Benjamini and Hochberg, 1995) procedure to control FDR. A gene was declared to have a differential expression if its adjusted *p*-value was lower than 0.05. The FPKM value (Fragments Per Kilobase of transcript per Million mapped reads) is used to estimate and compare gene expressions in eFP Browser. This normalization is based on the number of paired-end reads that mapped each gene taking into account the gene length and the library size.

### Comparison of RNA-seq data-set and hierarchical clustering

To be compared to our dataset, the raw RNA-seq reads published by Zhan et al. (2015) were retrieve from NCBI SRA (Leinonen et al., 2011) from Bioproject PRJNA265095 (runs SRR1633457 to SRR1633478). That represents 53 millions of pairs of length 2×100 bases for 22 samples. The reads from the two datasets were processed using the same pipeline: quality control was performed using FastQC version 0.11.7 (S. Andrew, http://www.bioinformatics.babraham.ac.uk/projects/fastqc/). Sequencing adapters were clipped using cutadapt v1.16 (Martin, 2011) sequencer artefacts were removed using FASTX Toolkit version 0.0.14 (http://hannonlab.cshl.edu/fastx_toolkit/), and custom Perl scripts were applied to trim regions of reads having an average Phred quality score (Ewing and Green, 1998) lower than 28 bases over a sliding window of 4 bases. We noticed some samples retrieved from SRA exhibited a high ribosomal RNA content. We built a maize rRNA database by comparing sequences from Silva (Quast et al., 2013) and RFAM (Kalvari et al., 2018) to the *Zea mays* B73 genome sequence; we then used this custom database to filter the RNA-seq reads with sortMeRNA version 2.1b (Kopylova et al., 2012). Reads shorter than 25 bases at the end of this processing, or with no mate, were discarded.

The genomic mapper HISAT2 v2.2.0 (Kim et al., 2015) was used to align read pairs against the *Zea mays* B73 genome sequence (AGP v4; Jiao et al., 2017), using the gene annotation version 4.40 provided as a GFF file (Wang et al., 2016); a first mapping pass was performed with the complete set of read pairs to discover unannotated splicing sites, before the per-sample mapping, with options “-k 10 --no-discordant –no-softclip” and allowing introns of length 40 to 150,000 bp. Mapped reads were counted by gene (not distinguishing isoforms) using FeatureCounts (Liao et al., 2014). The genome sequence and annotation file used was retrieved from the Gramene database (http://www.gramene.org/, release 51, in September 2016; Gupta et al., 2016).

Normalization, differential analysis and PCAs were performed with DESeq2 (Love et al., 2014) under R version 3.5.3 (R Development Core Team, 2005). In parallel, FPKM values (Fragments Per Kilobase of transcript per Million mapped reads) and confidence intervals were estimated using Cufflinks version 2.2.1 (Roberts et al., 2011) with options “--frag-bias-correct --multi-read-correct --max-multiread-fraction 1”.

### Functional annotation of *Zea mays* transcriptome, GO term enrichment analysis

The *Zea mays* B73 genome sequence v4 (Jiao et al., 2017) and the gene annotation v4.40 were used to predict transcript sequences using the gffread script from the Cufflinks package v2.2.1 (Trapnell et al., 2013). In each isoform sequence, the putative ORFs were identified using TransDecoder ((Haas et al., 2013) https://github.com/TransDecoder/TransDecoder/wiki), and the amino acid sequence was predicted. From 46 272 genes, 138 270 transcripts were predicted, leading to 149 699 amino acid sequences.

The predicted protein sequences were annotated for functional domains with InterProScan v5.27-66.0 ((Jones et al., 2014) using databases Pfam v31.0 ((Punta et al., 2012) and Panther 12.0 (Mi et al., 2013)). They were also compared to UniprotKB protein database version 2017_12 (2019). The complete Swiss-Prot database of curated proteins was used, (containing 41 689 plant sequences and 514 699 non-plant sequences) but only the plant subset of the non-curated database TrEMBL, containing 5 979 810 sequences). The comparison was carried out using WU-BlastP v2.0MP (Altschul et al., 1990) with parameters “W=3 Q=7 R=2 matrix=BLOSUM80 B=200 V=200 E=1e-6 hitdist=60 hspsepqmax=30 hspsepsmax=30 sump postsw”. Blast output was filtered using custom Perl scripts to keep only matches with log_10_ (e-value) no lower than 75% of the best log_10_ (e-value). GeneOntology terms (Ashburner et al., 2000) associated to matched proteins were retrieved from AmiGO (Carbon et al., 2009) with all their ancestors in the GO graph, using the SQL interface. For each *Zea mays* protein, we kept the GO terms associated to all its matched proteins or at least to 5 matched proteins. For each *Zea mays* gene, we merged the GO terms of all its isoforms.

For subsets of genes selected based on their expression pattern, we used our GO annotation to perform an enrichment analysis. The enrichment of a gene subset in a specific GO term is defined as:

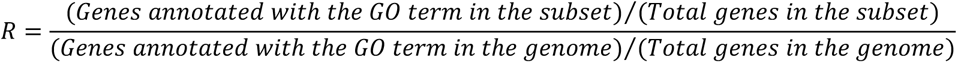

A hypergeometric test (R version 3.2.3; R Development Core Team (2005)) was applied to assess the significance of enrichment/depletion of each subset (Falcon and Gentleman, 2007; Pavlidis et al., 2004). Custom Perl scripts using GraphViz ((Ellson et al., 2001) https://graphviz.gitlab.io/) were used to browse the GeneOntology graph and identify enrichments or depletions that were both statistically significant and biologically relevant.

### Analysis of gene categories and orthology

Analysis of orthology to rice and Arabidopsis (Table 3) was based on Maize GDB annotations (https://www.maizegdb.org/, Andorf et al., 2016)). *Zein* genes were selected based on a previous gene list (Chen et al., 2014, 2017) and on Gramene database annotations (http://www.gramene.org/, Gupta et al., 2016)). The list of cell death associated genes was based on previously published lists (Arora et al., 2017; Fagundes et al., 2015). Heat maps were drawn with the online Heatmapper tool (http://www2.heatmapper.ca/, Babicki et al., 2016).

### Kernel fixation and *in situ* hybridization

Kernels were fixed in 4% of paraformaldehyde (pH7 adjusted with H_2_SO_4_) for 2 h under vacuum. For increased fixation efficiency, the two upper corners of the kernels were cut and vacuum was broken every 15 min. Kernels were dehydrated and included with Paraplast according to the protocol described by Jackson, 1991. Sections of 10-15 µm were cut with a HM355S microtome and attached on Adhesion Slides Superfrost Ultra plus^TM^ (ThermoFisher Scientific). RNA probes were amplified from genomic or cDNA (Supplemental Table 4) and labelled by digoxigenin (DIG) using the T7 reverse transcriptase kit of Promega, according to company instructions. RNA probes were then hydrolysed in carbonate buffer (120 mM Na_2_CO_3_, 80 mM NaHCO_3_) at 60°C for various times depending on the probe length (Supplemental Table 4) in order to obtain RNA fragments between 200 and 300 nucleotides (Jackson, 1991).

For the pre-hybridization of the sections, the protocol described by Jackson in 1991 was followed with some slight changes: pronase was replaced by proteinase K (1 µg.mL^-1^, ThermoFisher Scientific) in its buffer (100 mM Tris, 50 mM EDTA, pH8) and formaldehyde was replaced by paraformaldehyde as described above. For each slide, 1 µL of RNA probe was diluted in 74 µL of DIG easy Hyb buffer (Roche), denatured for 3 minutes at 80°C and dropped on a section that was immediately covered by a coverslip. Hybridization was carried out overnight at 50°C, in a hermetically closed box. Initial post hybridization treatments were carried out using gentle shaking as follows: 0.1X SSC buffer (from stock solution 20X SSC (3M NaCl, 300mM trisodium citrate, adjusted to pH7 with HCl)) and 0.5% SDS for 30 min at 50°C to remove the coverslips. Two baths of 1 h 30 in 2X SSC buffer mixed with 50% of formamide at 50°C and followed by 5 min in TBS buffer (400 mM NaCl, 0.1 mM Tris/HCl, pH7.5) at room temperature. Slides were then incubated in 0.5% blocking reagent solution (Roche) for 1h, followed by 30 min in TBS buffer with 1% BSA and 0.3% triton X100. Probes immunodetection was carried out in a wet chamber with 500 µL per slide of 0.225 U.mL^-1^ anti-DIG antibodies (Anti-Digoxigenin-AP, Fab fragments, Sigma-Aldrich) diluted in TBS with 1% BSA and 0.3% triton X100. After 1 h 30 of incubation, slides were washed 3 times 20 min in TBS buffer with 1% BSA, 0.3% triton and equilibrated in buffer 5 (100 mM Tris/HCl pH9.5, 100 mM NaCl, 50 mM MgCl_2_). Revelation was performed overnight in darkness in a buffer with 0.5 g.L^-1^ of nitroblue tetrazolium (NBT) and 0.2 g.L^-1^ of 5-Bromo-4-chloro-3-indolyl phosphate (BCIP). Slides were finally washed 4 times in water to stop the reaction and were optionally stained with calcofluor (fluorescent brightener 28, Sigma-Aldrich) and mounted in entellan (VWR). Pictures were taken either with VHX900F digital microscope (Keyence) or for magnification with Axio Imager 2 microscope (Zeiss).

### TUNEL staining

Fifteen DAP kernels were fixed in PFA, included in Paraplast and sectioned as described above. Paraplast was removed by successive baths in xylene (2x 5 min) and samples were then rehydrated through the following ethanol series: ethanol 100% (5 min), ethanol 95% (3 min), ethanol 70% (3 min), ethanol 50% (3 min), NaCl 0.85% in water (5 min) and Dulbecco’s Phosphate-Buffered Saline solution (PBS) (5 min). Sections were then permeabilized using proteinase K (1 µg/mL, ThermoFisher Scientific) for 10 min at 37°C and fixed again in PFA. Sections were washed in PBS and TUNEL staining was carried out with the ApoAlert DNA Fragmentation Assay Kit (Takara) according to manufacturer’s instructions. Sections were then counter-stained with propidium iodide (1 µg.ml^-1^ in PBS) for 15 min in darkness before being washed three times 5 min in water. Slides were mounted in Anti-fade Vectashield (Vector Laboratories). The fluorescein-dUTP incorporated at the free 3’-hydroxyl ends of fragmented DNA was excited at 520nm and propidium iodide at 620nm. Images were taken on a spinning disk microscope, with a CSU22 confocal head (Yokogawa) and an Ixon897 EMCCD camera (Andor) on a DMI4000 microscope (Leica).

### Data Deposition

RNA-Seq raw data were deposited in the international repository GEO (Gene Expression Omnibus, Edgar et al., 2002, http://www.ncbi.nlm.nih.gov/geo) under project ID GSE110060 (*Available when publication will be accepted*). RNA-seq data as FPKM values is available via the eFP Browser engine (http://bar.utoronto.ca/efp_maize/cgi-bin/efpWeb.cgi?dataSource=Maize_Kernel), which ‘paints’ the expression data onto images representing the samples used to generate the RNA-seq data.

## Supporting information

merged_sup_data

## Acknowledgements

We acknowledge Justin Berger, Patrice Bolland and Alexis Lacroix for maize culture, Isabelle Desbouchages and Hervé Leyral for buffer and media preparation, as well as Jérôme Laplaige, Marie-France Gérentes and Ghislaine Gendrot for technical assistance during samples dissections. We also thank Sophy Chamot and Frédérique Rozier for sharing protocols for *in-situ* hybridization. The sequencing platform (POPS-IPS2) benefits from the support of the LabEx Saclay Plant Sciences-SPS (ANR-10-LABX-0040-SPS). We acknowledge support by the INRA Plant Science and Breeding Division for the project SeedCom to TW. NMD was funded by a PhD fellowship from the Ministère de l’Enseignement Supérieur et de la Recherche. Part of this work has been refused once for funding by the French granting agency ANR.

## Author contributions

NMD and TW conceived and designed the experiments. TW performed samples dissections and RNA extractions, JC performed RNA-seq library preparation and sequencing, VB performed RNA-seq read processing and gene expression analysis, JJ performed bioinformatics to create the GO database and provide scripts to analyses the GO, as well as realized the comparison between published transcriptomes, AG and NDF performed TUNEL assay, NMD performed all other remaining experiments. EE, AP and NJP contributed to the RNA-seq data accessibility via the eFP Browser engine. NMD, PMR and TW analysed the data. NMD prepared tables and figures. NMD, GI, PMR and TW wrote the manuscript. TW was involved in project management and obtained funding.

## Supplemental Tables

**Supplemental Table 1:** Number of kernels used for each of the four biological replicates. For each replicate, the material comes from two independent 13 day old maize ears.

|  | Embryo (Emb) | Endosperm (End) | Pericarp (Per) | Scutellar Aleurone Layer (SAL) | Apical Scutellum (AS) | Endosperm Adjacent to Scutellum (EAS) |
| --- | --- | --- | --- | --- | --- | --- |
| Rep-1 | 60 | 4 | 4 | 21 | 50 | 14 |
| Rep-2 | 50 | 4 | 4 | 15 | 72 | 15 |
| Rep-3 | 50 | 4 | 4 | 22 | 84 | 15 |
| Rep-4 | 50 | 4 | 4 | 19 | 66 | 16 |

**Supplemental Table 2:** Number of genes differentially expressed between a sub compartment and its compartment of origin. FDR = false discovery rate, DEGs = differentially expressed genes, AS = apical scutellum, Emb = embryo, End = endosperm, SAL = scutellar aleurone Layer, EAS = endosperm adjacent to scutellum.

|  | AS vs Emb | EAS vs End | SAL vs End |
| --- | --- | --- | --- |
| Number of genes with a $FDR \leq 0.05$ in the comparison | 11646 | 13161 | 16772 |
| Number of DEGs | 682 | 1498 | 2975 |
| Number of DEGs upregulated in the sub-compartment | 82 | 485 | 1995 |
| Number of DEGs downregulated in the sub-compartment | 600 | 1013 | 980 |

**Supplemental Table 3:** Mean expression values and gene IDs of genes selected for *in situ* hybridization.

| Name | Gene id (AGPv4) | Mean SAL | Mean Emb | Mean End | Mean EAS | Mean Per | Mean AS |
| --- | --- | --- | --- | --- | --- | --- | --- |
| *SWEET14a* | Zm00001d007365 | 562 | 50 | 666 | 10876 | 22 | 36 |
| *SWEET15a* | Zm00001d050577 | 3105 | 21 | 1935 | 31147 | 200 | 10 |
| *UMAMIT_EAS1* | Zm00001d009063 | 3,25 | 0,5 | 45,5 | 467 | 3 | 0 |
| *PEPB11* | Zm00001d037439 | 4586 | 24 | 3023 | 41507 | 265 | 13 |
| *unknown* | Zm00001d017285 | 668 | 3 | 468 | 7271 | 85 | 2 |
| *SCL49* | Zm00001d014983 | 692 | 449 | 2327 | 39382 | 4587 | 361 |
| *ZmNAC124* | Zm00001d046126 | 0 | 482 | 0 | 0 | 0 | 0,5 |

**Supplemental Table 4:**
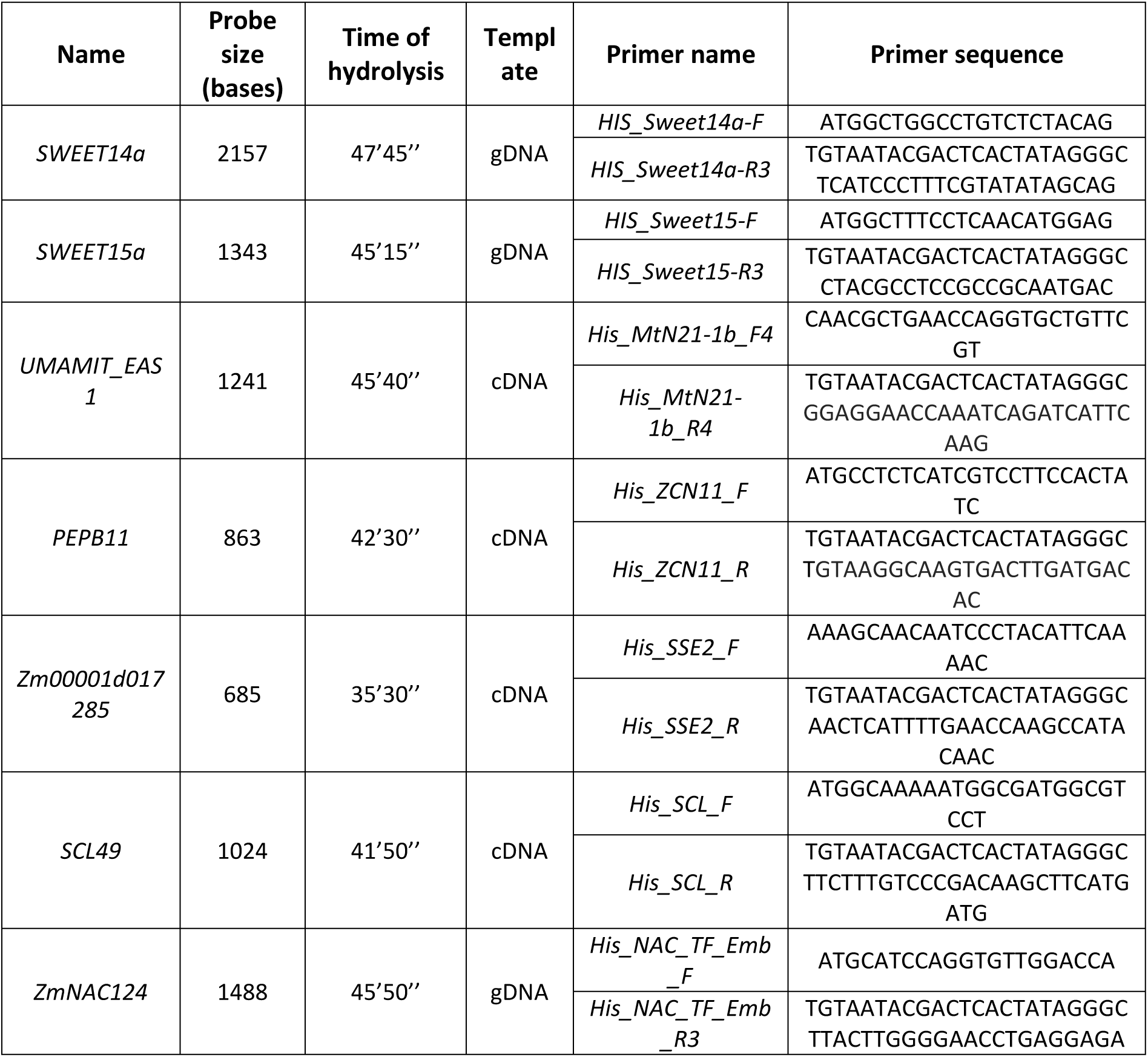
Primers used in this study and conditions for RNA probes synthesis.

**Supplemental Data Set 1:** number of normalized read counts per gene annotated in the AGP v4 version of the B73 maize genome. Read counts were normalized using the trimmed mean of M-value (TMM) method (library size normalization).

**Supplemental Data Set 2:** Pairwise comparison of gene expression levels between the tissues. The 15 different comparisons are presented on different sheets. The genes with a statistical difference supported by a FDR < 0.05 are listed. Among them, the DEGs (*i.e.* genes with a log_2_(fold change) ≥ 2) are highlighted in yellow.

## Bibliography

Altschul, S.F., Gish, W., Miller, W., Myers, E.W., and Lipman, D.J. (1990). Basic local alignment search tool. J. Mol. Biol. 215, 403–410.

Anders, S., Pyl, P.T., and Huber, W. (2015). HTSeq--a Python framework to work with high-throughput sequencing data. Bioinforma. Oxf. Engl. 31, 166–169.

Andorf, C.M., Cannon, E.K., Portwood, J.L., Gardiner, J.M., Harper, L.C., Schaeffer, M.L., Braun, B.L., Campbell, D.A., Vinnakota, A.G., Sribalusu, V.V., et al. (2016). MaizeGDB update: new tools, data and interface for the maize model organism database. Nucleic Acids Res. 44, D1195–D1201.

Arora, K., Panda, K.K., Mittal, S., Mallikarjuna, M.G., Rao, A.R., Dash, P.K., and Thirunavukkarasu, N. (2017). RNAseq revealed the important gene pathways controlling adaptive mechanisms under waterlogged stress in maize. Sci. Rep. 7.

Ashburner, M., Ball, C.A., Blake, J.A., Botstein, D., Butler, H., Cherry, J.M., Davis, A.P., Dolinski, K., Dwight, S.S., Eppig, J.T., et al. (2000). Gene Ontology: tool for the unification of biology. Nat. Genet. 25, 25–29.

Babicki, S., Arndt, D., Marcu, A., Liang, Y., Grant, J.R., Maciejewski, A., and Wishart, D.S. (2016). Heatmapper: web-enabled heat mapping for all. Nucleic Acids Res. 44, W147–153.

Belmonte, M.F., Kirkbride, R.C., Stone, S.L., Pelletier, J.M., Bui, A.Q., Yeung, E.C., Hashimoto, M., Fei, J., Harada, C.M., Munoz, M.D., et al. (2013). Comprehensive developmental profiles of gene activity in regions and subregions of the Arabidopsis seed. Proc. Natl. Acad. Sci. U. S. A. 110, E435–E444.

Benjamini, Y., and Hochberg, Y. (1995). Controlling the False Discovery Rate: A Practical and Powerful Approach to Multiple Testing. J. R. Stat. Soc. Ser. B Methodol. 57, 289–300.

Berger, F. (1999). Endosperm development. Curr. Opin. Plant Biol. 2, 28–32.

Berger, F. (2003). Endosperm: the crossroad of seed development. Curr. Opin. Plant Biol. 6, 42–50.

Bezrutczyk, M., Hartwig, T., Horschman, M., Char, S.N., Yang, J., Yang, B., Frommer, W.B., and Sosso, D. (2018). Impaired phloem loading in zmsweet13a,b,c sucrose transporter triple knock-out mutants in Zea mays. New Phytol. 218, 594–603.

Bommert, P., and Werr, W. (2001). Gene expression patterns in the maize caryopsis: clues to decisions in embryo and endosperm development. Gene 271, 131–142.

Bourgon, R., Gentleman, R., and Huber, W. (2010). Independent filtering increases detection power for high-throughput experiments. Proc. Natl. Acad. Sci. 107, 9546–9551.

Cai, G., Faleri, C., Del Casino, C., Hueros, G., Thompson, R.D., and Cresti, M. (2002). Subcellular localisation of BETL-1, -2 and -4 in Zea mays L. endosperm. Sex. Plant Reprod. 15, 85–98.

Carbon, S., Ireland, A., Mungall, C.J., Shu, S., Marshall, B., and Lewis, S. (2009). AmiGO: online access to ontology and annotation data. Bioinformatics 25, 288–289.

Chen, J., Zeng, B., Zhang, M., Xie, S., Wang, G., Hauck, A., and Lai, J. (2014). Dynamic Transcriptome Landscape of Maize Embryo and Endosperm Development. Plant Physiol. 166, 252–264.

Chen, L.-Q., Qu, X.-Q., Hou, B.-H., Sosso, D., Osorio, S., Fernie, A.R., and Frommer, W.B. (2012). Sucrose efflux mediated by SWEET proteins as a key step for phloem transport. Science 335, 207–211.

Chen, X., Feng, F., Qi, W., Xu, L., Yao, D., Wang, Q., and Song, R. (2017). Dek35 Encodes a PPR Protein that Affects cis-Splicing of Mitochondrial nad4 Intron 1 and Seed Development in Maize. Mol. Plant 10, 427–441.

Cheng, W.H., Taliercio, E.W., and Chourey, P.S. (1996). The Miniature1 seed locus of maize encodes a cell wall invertase required for normal development of endosperm and maternal cells in the pedicel. Plant Cell 8, 971–983.

Chourey, P.S., and Hueros, G. (2017). The basal endosperm transfer layer (BETL): Gateway to the maize kernel. In Maize Kernel Development, (Larkins BA), pp. 56–67.

Davis, R., Smith, J., and Cobb, B. (1990). A Light and Electron-Microscope Investigation of the Transfer Cell Region of Maize Caryopses. Can. J. Bot.-Rev. Can. Bot. 68, 471–479.

Diboll, A., and Larson, D. (1966). An electron microscopic study of the mature megagametophyte in Zea mays. Am. J. Bot. 391–402.

Doll, N.M., Depège-Fargeix, N., Rogowsky, P.M., and Widiez, T. (2017). Signaling in Early Maize Kernel Development. Mol. Plant 10, 375–388.

Downs, G.S., Bi, Y.-M., Colasanti, J., Wu, W., Chen, X., Zhu, T., Rothstein, S.J., and Lukens, L.N. (2013). A Developmental Transcriptional Network for Maize Defines Coexpression Modules. Plant Physiol. 161, 1830–1843.

Dumas, C., and Rogowsky, P. (2008). Fertilization and early seed formation. C. R. Biol. 331, 715–725.

Edgar, R., Domrachev, M., and Lash, A.E. (2002). Gene Expression Omnibus: NCBI gene expression and hybridization array data repository. Nucleic Acids Res. 30, 207–210.

Ellson, J., Gansner, E., Koutsofios, L., North, S., Woodhull, G., Description, S., and Technologies, L. (2001). Graphviz — open source graph drawing tools. In Lecture Notes in Computer Science, (Springer-Verlag), pp. 483–484.

Ewing, B., and Green, P. (1998). Base-calling of automated sequencer traces using phred. II. Error probabilities. Genome Res. 8, 186–194.

Fagundes, D., Bohn, B., Cabreira, C., Leipelt, F., Dias, N., Bodanese-Zanettini, M.H., and Cagliari, A. (2015). Caspases in plants: metacaspase gene family in plant stress responses. Funct. Integr. Genomics 15, 639–649.

Falcon, S., and Gentleman, R. (2007). Using GOstats to test gene lists for GO term association. Bioinforma. Oxf. Engl. 23, 257–258.

Feng, F., Qi, W., Lv, Y., Yan, S., Xu, L., Yang, W., Yuan, Y., Chen, Y., Zhao, H., and Song, R. (2018). OPAQUE11 Is a Central Hub of the Regulatory Network for Maize Endosperm Development and Nutrient Metabolism. Plant Cell 30, 375–396.

Fourquin, C., Beauzamy, L., Chamot, S., Creff, A., Goodrich, J., Boudaoud, A., and Ingram, G. (2016). Mechanical stress mediated by both endosperm softening and embryo growth underlies endosperm elimination in Arabidopsis seeds. Dev. Camb. Engl. 143, 3300–3305.

Gagnot, S., Tamby, J.-P., Martin-Magniette, M.-L., Bitton, F., Taconnat, L., Balzergue, S., Aubourg, S., Renou, J.-P., Lecharny, A., and Brunaud, V. (2008). CATdb: a public access to Arabidopsis transcriptome data from the URGV-CATMA platform. Nucleic Acids Res. 36, D986–990.

Galluzzi, L., Bravo-San Pedro, J.M., Vitale, I., Aaronson, S.A., Abrams, J.M., Adam, D., Alnemri, E.S., Altucci, L., Andrews, D., Annicchiarico-Petruzzelli, M., et al. (2015). Essential versus accessory aspects of cell death: recommendations of the NCCD 2015. Cell Death Differ. 22, 58–73.

Giuliani, C., Consonni, G., Gavazzi, G., Colombo, M., and Dolfini, S. (2002). Programmed cell death during embryogenesis in maize. Ann. Bot. 90, 287–292.

Gómez, E., Royo, J., Guo, Y., Thompson, R., and Hueros, G. (2002). Establishment of Cereal Endosperm Expression Domains Identification and Properties of a Maize Transfer Cell–Specific Transcription Factor, ZmMRP-1. Plant Cell 14, 599–610.

Gomez, E., Royo, J., Muniz, L.M., Sellam, O., Paul, W., Gerentes, D., Barrero, C., Lopez, M., Perez, P., and Hueros, G. (2009). The Maize Transcription Factor Myb-Related Protein-1 Is a Key Regulator of the Differentiation of Transfer Cells. Plant Cell 21, 2022–2035.

Gontarek, B.C., and Becraft, P.W. (2017). Aleurone. In Maize Kernel Development, B. Larkins, ed. (Wallingford: CABI), pp. 68–80.

Grimault, A., Gendrot, G., Chamot, S., Widiez, T., Rabille, H., Gerentes, M.-F., Creff, A., Thevenin, J., Dubreucq, B., Ingram, G.C., et al. (2015). ZmZHOUPI, an endosperm-specific basic helix-loop-helix transcription factor involved in maize seed development. Plant J. 84, 574–586.

Gupta, P., Naithani, S., Tello-Ruiz, M.K., Chougule, K., D’Eustachio, P., Fabregat, A., Jiao, Y., Keays, M., Lee, Y.K., Kumari, S., et al. (2016). Gramene Database: Navigating Plant Comparative Genomics Resources. Curr. Plant Biol. 7–8, 10.

Gutiérrez-Marcos, J.F., Costa, L.M., Biderre-Petit, C., Khbaya, B., O’Sullivan, D.M., Wormald, M., Perez, P., and Dickinson, H.G. (2004). maternally expressed gene1 Is a Novel Maize Endosperm Transfer Cell–Specific Gene with a Maternal Parent-of-Origin Pattern of Expression. Plant Cell 16, 1288–1301.

Haas, B.J., Papanicolaou, A., Yassour, M., Grabherr, M., Blood, P.D., Bowden, J., Couger, M.B., Eccles, D., Li, B., Lieber, M., et al. (2013). *De novo* transcript sequence reconstruction from RNA-seq using the Trinity platform for reference generation and analysis. Nat. Protoc. 8, 1494–1512.

Heckel, T., Werner, K., Sheridan, W.F., Dumas, C., and Rogowsky, P.M. (1999). Novel phenotypes and developmental arrest in early embryo specific mutants of maize. Planta 210, 1–8.

Hueros, G., Royo, J., Maitz, M., Salamini, F., and Thompson, R.D. (1999a). Evidence for factors regulating transfer cell-specific expression in maize endosperm. Plant Mol. Biol. 41, 403–414.

Hueros, G., Gomez, E., Cheikh, N., Edwards, J., Weldon, M., Salamini, F., and Thompson, R.D. (1999b). Identification of a Promoter Sequence from the BETL1Gene Cluster Able to Confer Transfer-Cell-Specific Expression in Transgenic Maize. Plant Physiol. 121, 1143–1152.

Ingram, G., and Gutierrez-Marcos, J. (2015). Peptide signalling during angiosperm seed development. J. Exp. Bot. 66, 5151–5159.

Ingram, G.C., Boisnard-Lorig, C., Dumas, C., and Rogowsky, P.M. (2000). Expression patterns of genes encoding HD-ZipIV homeo domain proteins define specific domains in maize embryos and meristems. Plant J. Cell Mol. Biol. 22, 401–414.

Jackson, D. (1991). In-situ hybridization in plants. In Molecular Plant Pathology: A Practical Approach, (Bowles DJ), pp. 163–174.

Jestin, L., Ravel, C., Auroy, S., Laubin, B., Perretant, M.-R., Pont, C., and Charmet, G. (2008). Inheritance of the number and thickness of cell layers in barley aleurone tissue (Hordeum vulgare L.): an approach using F2-F3 progeny. Theor. Appl. Genet. 116, 991–1002.

Jiao, Y., Peluso, P., Shi, J., Liang, T., Stitzer, M.C., Wang, B., Campbell, M.S., Stein, J.C., Wei, X., Chin, C.-S., et al. (2017). Improved maize reference genome with single-molecule technologies. Nature 546, 524–527.

Jones, P., Binns, D., Chang, H.-Y., Fraser, M., Li, W., McAnulla, C., McWilliam, H., Maslen, J., Mitchell, A., Nuka, G., et al. (2014). InterProScan 5: genome-scale protein function classification. Bioinformatics 30, 1236.

Kalvari, I., Argasinska, J., Quinones-Olvera, N., Nawrocki, E.P., Rivas, E., Eddy, S.R., Bateman, A., Finn, R.D., and Petrov, A.I. (2018). Rfam 13.0: shifting to a genome-centric resource for non-coding RNA families. Nucleic Acids Res. 46, D335–D342.

Kang, B.-H., Xiong, Y., Williams, D.S., Pozueta-Romero, D., and Chourey, P.S. (2009). Miniature1-Encoded Cell Wall Invertase Is Essential for Assembly and Function of Wall-in-Growth in the Maize Endosperm Transfer Cell. Plant Physiol. 151, 1366–1376.

Kiesselbach, T.A. (1949). The Structure and Reproduction of Corn (CSHL Press).

Kiesselbach, T.A., and Walker, E.R. (1952). Structure of Certain Specialized Tissues in the Kernel of Corn. Am. J. Bot. 39, 561–569.

Kim, D., Langmead, B., and Salzberg, S.L. (2015). HISAT: a fast spliced aligner with low memory requirements. Nat. Methods 12, 357–360.

Kladnik, A., Chamusco, K., Dermastia, M., and Chourey, P. (2004). Evidence of programmed cell death in post-phloem transport cells of the maternal pedicel tissue in developing caryopsis of maize. Plant Physiol. 136, 3572–3581.

Kopylova, E., Noé, L., and Touzet, H. (2012). Kopylova E, Noe L, Touzet H.. SortMeRNA: Fast and accurate filtering of ribosomal RNAs in metatranscriptomic data. Bioinformatics 28: 3211-3217. Bioinforma. Oxf. Engl. 28, 3211–3217.

Labat-Moleur, F., Guillermet, C., Lorimier, P., Robert, C., Lantuejoul, S., Brambilla, E., and Negoescu, A. (1998). TUNEL Apoptotic Cell Detection in Tissue Sections: Critical Evaluation and Improvement. J. Histochem. Cytochem. 46, 327–334.

Langmead, B., and Salzberg, S.L. (2012). Fast gapped-read alignment with Bowtie 2. Nat. Methods 9, 357–359.

Le, B.H., Cheng, C., Bui, A.Q., Wagmaister, J.A., Henry, K.F., Pelletier, J., Kwong, L., Belmonte, M., Kirkbride, R., Horvath, S., et al. (2010). Global analysis of gene activity during Arabidopsis seed development and identification of seed-specific transcription factors. Proc. Natl. Acad. Sci. 107, 8063–8070.

Leinonen, R., Sugawara, H., Shumway, M., and International Nucleotide Sequence Database Collaboration (2011). The sequence read archive. Nucleic Acids Res. 39, D19–21.

Leroux, B.M., Goodyke, A.J., Schumacher, K.I., Abbott, C.P., Clore, A.M., Yadegari, R., Larkins, B.A., and Dannenhoffer, J.M. (2014). Maize early endosperm growth and development: From fertilization through cell type differentiation. Am. J. Bot. 101, 1259–1274.

Li, G., Wang, D., Yang, R., Logan, K., Chen, H., Zhang, S., Skaggs, M.I., Lloyd, A., Burnett, W.J., Laurie, J.D., et al. (2014). Temporal patterns of gene expression in developing maize endosperm identified through transcriptome sequencing. Proc. Natl. Acad. Sci. U. S. A. 111, 7582–7587.

Liao, Y., Smyth, G.K., and Shi, W. (2014). featureCounts: an efficient general purpose program for assigning sequence reads to genomic features. Bioinforma. Oxf. Engl. 30, 923–930.

Lopes, M.A., and Larkins, B.A. (1993). Endosperm origin, development, and function. Plant Cell 5, 1383–1399.

Love, M.I., Huber, W., and Anders, S. (2014). Moderated estimation of fold change and dispersion for RNA-seq data with DESeq2. Genome Biol. 15, 550.

Lowe, J., and Nelson, O. (1946). Miniature Seed - a Study in the Development of a Defective Caryopsis in Maize. Genetics 31, 525-.

Lu, X., Chen, D., Shu, D., Zhang, Z., Wang, W., Klukas, C., Chen, L., Fan, Y., Chen, M., and Zhang, C. (2013). The Differential Transcription Network between Embryo and Endosperm in the Early Developing Maize Seed(1[C][W][OA]). Plant Physiol. 162, 440–455.

Martin, M. (2011). Cutadapt removes adapter sequences from high-throughput sequencing reads. EMBnet.Journal 17, 10–12.

McCarthy, D.J., Chen, Y., and Smyth, G.K. (2012). Differential expression analysis of multifactor RNA-Seq experiments with respect to biological variation. Nucleic Acids Res. 40, 4288–4297.

Meng, D., Zhao, J., Zhao, C., Luo, H., Xie, M., Liu, R., Lai, J., Zhang, X., and Jin, W. (2018). Sequential gene activation and gene imprinting during early embryo development in maize. Plant J. Cell Mol. Biol. 93, 445–459.

Mi, H., Muruganujan, A., and Thomas, P.D. (2013). PANTHER in 2013: modeling the evolution of gene function, and other gene attributes, in the context of phylogenetic trees. Nucleic Acids Res. 41, D377–386.

Miller, M., and Chourey, P. (1992). The Maize Invertase-Deficient Miniature-1 Seed Mutation Is Associated with Aberrant Pedicel and Endosperm Development. Plant Cell 4, 297–305.

Mimura, M., Kudo, T., Wu, S., McCarty, D.R., and Suzuki, M. (2018). Autonomous and nonautonomous functions of the maize Shohai1 gene, encoding a RWP-RK putative transcription factor, in regulation of embryo and endosperm development. Plant J. Cell Mol. Biol.

Müller, B., Fastner, A., Karmann, J., Mansch, V., Hoffmann, T., Schwab, W., Suter-Grotemeyer, M., Rentsch, D., Truernit, E., Ladwig, F., et al. (2015). Amino Acid Export in Developing Arabidopsis Seeds Depends on UmamiT Facilitators. Curr. Biol. 25, 3126–3131.

Nelson, O., and Pan, D. (1995). Starch Synthesis in Maize Endosperms. Annu. Rev. Plant Physiol. Plant Mol. Biol. 46, 475–496.

Norholm, M.H.H., Nour-Eldin, H.H., Brodersen, P., Mundy, J., and Halkier, B.A. (2006). Expression of the Arabidopsis high-affinity hexose transporter STP13 correlates with programmed cell death. FEBS Lett. 580, 2381–2387.

Nowack, M.K., Ungru, A., Bjerkan, K.N., Grini, P.E., and Schnittger, A. (2010). Reproductive cross-talk: seed development in flowering plants. Biochem. Soc. Trans. 38, 604–612.

Olsen, O.-A. (2001). ENDOSPERM DEVELOPMENT: Cellularization and Cell Fate Specification. Annu. Rev. Plant Physiol. Plant Mol. Biol. 52, 233–267.

Olsen, O.A. (2004a). Dynamics of maize aleurone cell formation: The “surface-”rule. Maydica 49, 37–40.

Olsen, O.-A. (2004b). Nuclear Endosperm Development in Cereals and Arabidopsis thaliana. Plant Cell 16, S214–S227.

Olvera-Carrillo, Y., Van Bel, M., Van Hautegem, T., Fendrych, M., Huysmans, M., Simaskova, M., van Durme, M., Buscaill, P., Rivas, S., S Coll, N., et al. (2015). A Conserved Core of Programmed Cell Death Indicator Genes Discriminates Developmentally and Environmentally Induced Programmed Cell Death in Plants. Plant Physiol. 169, 2684–2699.

OpsahlFerstad, H.G., LeDeunff, E., Dumas, C., and Rogowsky, P.M. (1997). ZmEsr, a novel endosperm-specific gene expressed in a restricted region around the maize embryo. Plant J. 12, 235–246.

Pavlidis, P., Qin, J., Arango, V., Mann, J.J., and Sibille, E. (2004). Using the gene ontology for microarray data mining: a comparison of methods and application to age effects in human prefrontal cortex. Neurochem. Res. 29, 1213–1222.

Porter, G.A., Knievel, D.P., and Shannon, J.C. (1987). Assimilate Unloading from Maize (Zea mays L.) Pedicel Tissues : II. Effects of Chemical Agents on Sugar, Amino Acid, and C-Assimilate Unloading. Plant Physiol. 85, 558–565.

Punta, M., Coggill, P.C., Eberhardt, R.Y., Mistry, J., Tate, J., Boursnell, C., Pang, N., Forslund, K., Ceric, G., Clements, J., et al. (2012). The Pfam protein families database. Nucleic Acids Res. 40, D290–301.

Qu, J., Ma, C., Feng, J., Xu, S., Wang, L., Li, F., Li, Y., Zhang, R., Zhang, X., Xue, J., et al. (2016). Transcriptome Dynamics during Maize Endosperm Development. PloS One 11, e0163814.

Quast, C., Pruesse, E., Yilmaz, P., Gerken, J., Schweer, T., Yarza, P., Peplies, J., and Glöckner, F.O. (2013). The SILVA ribosomal RNA gene database project: improved data processing and web-based tools. Nucleic Acids Res. 41, D590–596.

R Development Core Team (2005). A language and environment for statistical computing, reference index version 2.2.1.

Randolph, L.F. (1936). Developmental morphology of the caryopsis in maize ([U.S. Dept. of Agriculture]).

Rigaill, G., Balzergue, S., Brunaud, V., Blondet, E., Rau, A., Rogier, O., Caius, J., Maugis-Rabusseau, C., Soubigou-Taconnat, L., Aubourg, S., et al. (2018). Synthetic data sets for the identification of key ingredients for RNA-seq differential analysis. Brief. Bioinform. 19, 65–76.

Roberts, A., Trapnell, C., Donaghey, J., Rinn, J.L., and Pachter, L. (2011). Improving RNA-Seq expression estimates by correcting for fragment bias. Genome Biol. 12, R22.

Rousseau, D., Widiez, T., Di Tommaso, S., Rositi, H., Adrien, J., Maire, E., Langer, M., Olivier, C., Peyrin, F., and Rogowsky, P. (2015). Fast virtual histology using X-ray in-line phase tomography: application to the 3D anatomy of maize developing seeds. Plant Methods 11, 55.

Sabelli, P.A., and Larkins, B.A. (2009). The Development of Endosperm in Grasses. Plant Physiol. 149, 14–26.

Schmidt, R.J., Burr, F.A., Aukerman, M.J., and Burr, B. (1990). Maize regulatory gene opaque-2 encodes a protein with a “leucine-zipper” motif that binds to zein DNA. Proc. Natl. Acad. Sci. 87, 46–50.

Schon, M.A., and Nodine, M.D. (2017). Widespread Contamination of Arabidopsis Embryo and Endosperm Transcriptome Data Sets. Plant Cell 29, 608–617.

Sekhon, R.S., Lin, H., Childs, K.L., Hansey, C.N., Buell, C.R., de Leon, N., and Kaeppler, S.M. (2011). Genome-wide atlas of transcription during maize development. Plant J. Cell Mol. Biol. 66, 553–563.

Sosso, D., Canut, M., Gendrot, G., Dedieu, A., Chambrier, P., Barkan, A., Consonni, G., and Rogowsky, P.M. (2012). PPR8522 encodes a chloroplast-targeted pentatricopeptide repeat protein necessary for maize embryogenesis and vegetative development. J. Exp. Bot. 63, 5843–5857.

Sosso, D., Luo, D., Li, Q.-B., Sasse, J., Yang, J., Gendrot, G., Suzuki, M., Koch, K.E., McCarty, D.R., Chourey, P.S., et al. (2015). Seed filling in domesticated maize and rice depends on SWEET-mediated hexose transport. Nat. Genet. 47, 1489–1493.

Sreenivasulu, N., and Wobus, U. (2013). Seed-development programs: a systems biology-based comparison between dicots and monocots. Annu. Rev. Plant Biol. 64, 189–217.

Suzuki, M., Ketterling, M.G., Li, Q.-B., and McCarty, D.R. (2003). Viviparous1 alters global gene expression patterns through regulation of abscisic acid signaling. Plant Physiol. 132, 1664–1677.

Trapnell, C., Hendrickson, D.G., Sauvageau, M., Goff, L., Rinn, J.L., and Pachter, L. (2013). Differential analysis of gene regulation at transcript resolution with RNA-seq. Nat. Biotechnol. 31, 46–53.

Upadhyay, N., Kar, D., Deepak Mahajan, B., Nanda, S., Rahiman, R., Panchakshari, N., Bhagavatula, L., and Datta, S. The multitasking abilities of MATE transporters in plants. J. Exp. Bot.

Van Lammeren, A.A.M. van (1987). Embryogenesis in Zea mays L. : a structural approach to maize caryopsis development in vivo and in vitro.

Vernoud, V., Hajduch, M., Khaled, A.-S., Depege, N., and Rogowsky, P.M. (2005). Maize Embryogenesis. Maydica 50, 469–483.

Wang, B., Tseng, E., Regulski, M., Clark, T.A., Hon, T., Jiao, Y., Lu, Z., Olson, A., Stein, J.C., and Ware, D. (2016). Unveiling the complexity of the maize transcriptome by single-molecule long-read sequencing. Nat. Commun. 7, 11708.

Widiez, T., Ingram, G.C., and Gutiérrez-Marcos, J.F. (2017). Embryo-endosperm-sporophyte interactions in maize seeds. In Maize Kernel Development, B. Larkins, ed. (Wallingford: CABI), pp. 95–107.

Woo, Y.-M., Hu, D.W.-N., Larkins, B.A., and Jung, R. (2001). Genomics Analysis of Genes Expressed in Maize Endosperm Identifies Novel Seed Proteins and Clarifies Patterns of Zein Gene Expression. Plant Cell 13, 2297–2318.

Wu, Y., and Messing, J. (2014). Proteome balancing of the maize seed for higher nutritional value. Front. Plant Sci. 5, 240.

Yi, F., Gu, W., Chen, J., Song, N., Gao, X., Zhang, X., Zhou, Y., Ma, X., Song, W., Zhao, H., et al. (2019). High-temporal-resolution Transcriptome Landscape of Early Maize Seed Development. Plant Cell tpc.00961.2018.

Young, T.E., and Gallie, D.R. (2000). Programmed cell death during endosperm development. Plant Mol. Biol. 44, 283–301.

Zhan, J., Thakare, D., Ma, C., Lloyd, A., Nixon, N.M., Arakaki, A.M., Burnett, W.J., Logan, K.O., Wang, D., Wang, X., et al. (2015). RNA Sequencing of Laser-Capture Microdissected Compartments of the Maize Kernel Identifies Regulatory Modules Associated with Endosperm Cell Differentiation. Plant Cell 27, 513–531.

Zhan, J., Dannenhoffer, J.M., and Yadegari, R. (2017). Endosperm development and cell specialization. In Maize Kernel Development, B. Larkins, ed. (Wallingford: CABI), pp. 28–43.

Zhang, S., Wong, L., Meng, L., and Lemaux, P.G. (2002). Similarity of expression patterns of knotted1 and ZmLEC1 during somatic and zygotic embryogenesis in maize (Zea mays L.). Planta 215, 191–194.

Zheng, Y., and Wang, Z. (2014). Differentiation mechanism and function of the cereal aleurone cells and hormone effects on them. Plant Cell Rep. 33, 1779–1787.

Zheng, Y., and Wang, Z. (2015). The cereal starch endosperm development and its relationship with other endosperm tissues and embryo. Protoplasma 252, 33–40.

Zimmermann, R., and Werr, W. (2005). Pattern Formation in the Monocot Embryo as Revealed by NAMand CUC3 Orthologues from Zea mays L. Plant Mol. Biol. 58, 669–685.

(2019). UniProt: a worldwide hub of protein knowledge. Nucleic Acids Res. 47, D506–D515.

