## Supplementary material for "Transcriptomics at maize embryo/endosperm interfaces identify a novel transcriptionally distinct endosperm sub-domain adjacent to the embryo scutellum (EAS)": merged_sup_data

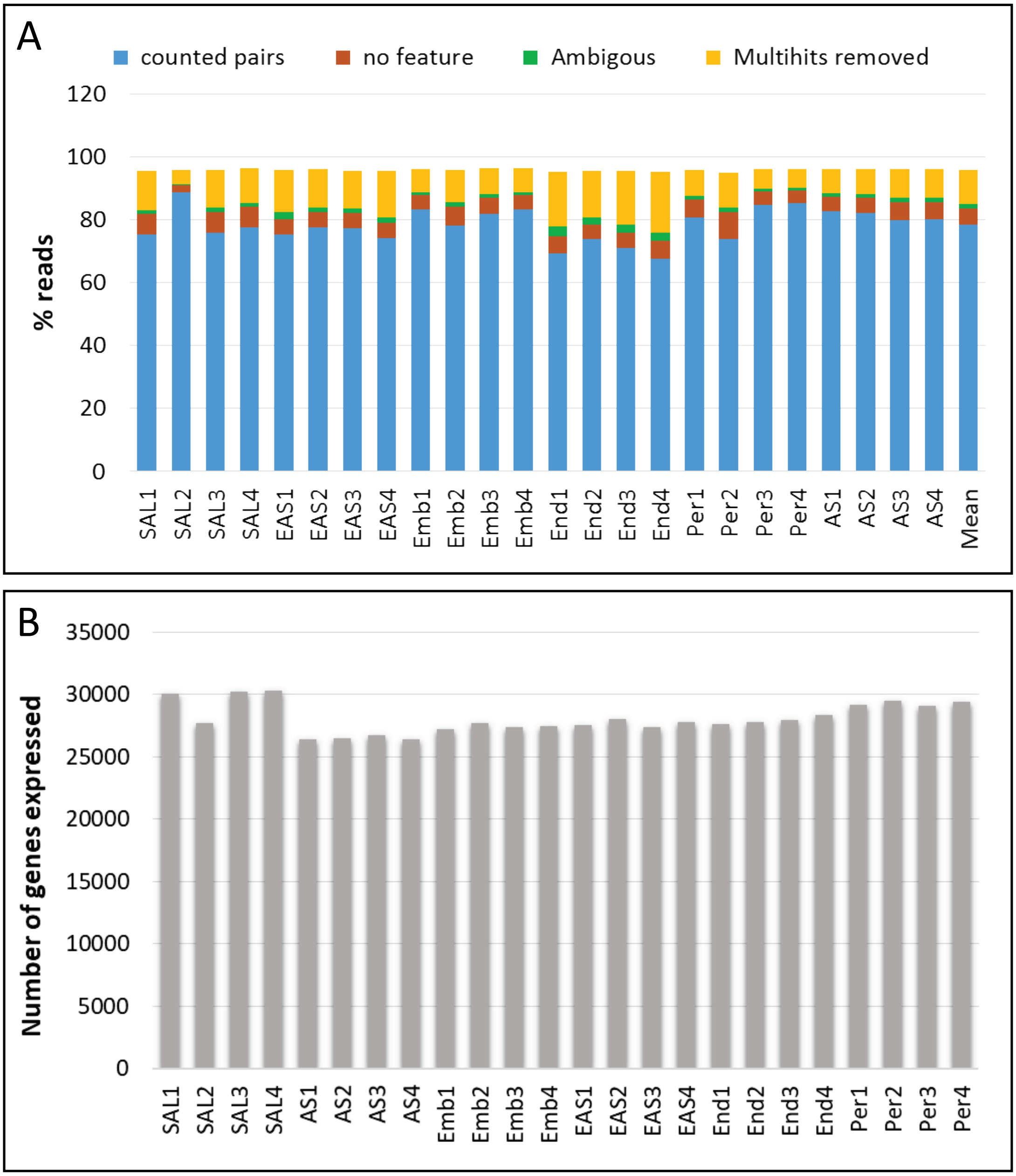


**Supplemental Figure 1**. **(A)** Proportion of reads mapped and filtered per sample. **(B)** Number of genes expressed with a normalized read counts over 1 for every sample.

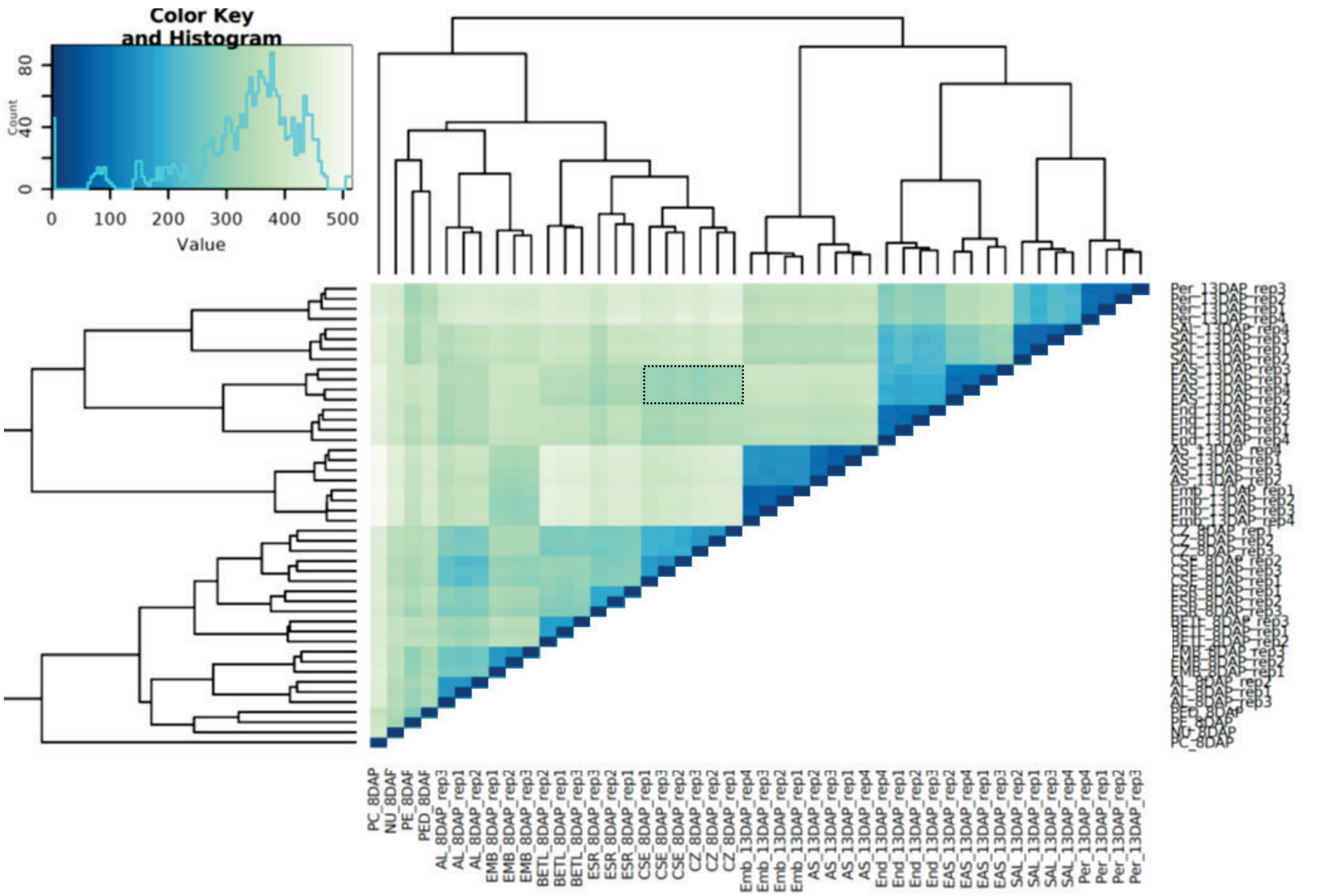


**Supplemental Figure 2.** Comparison between the transcriptomic data generated at 13 DAP in this study, and those at 8 DAP published in Zhan et al, 2015. Rectangle with dotted line shows that EAS samples transcriptome is closer to.

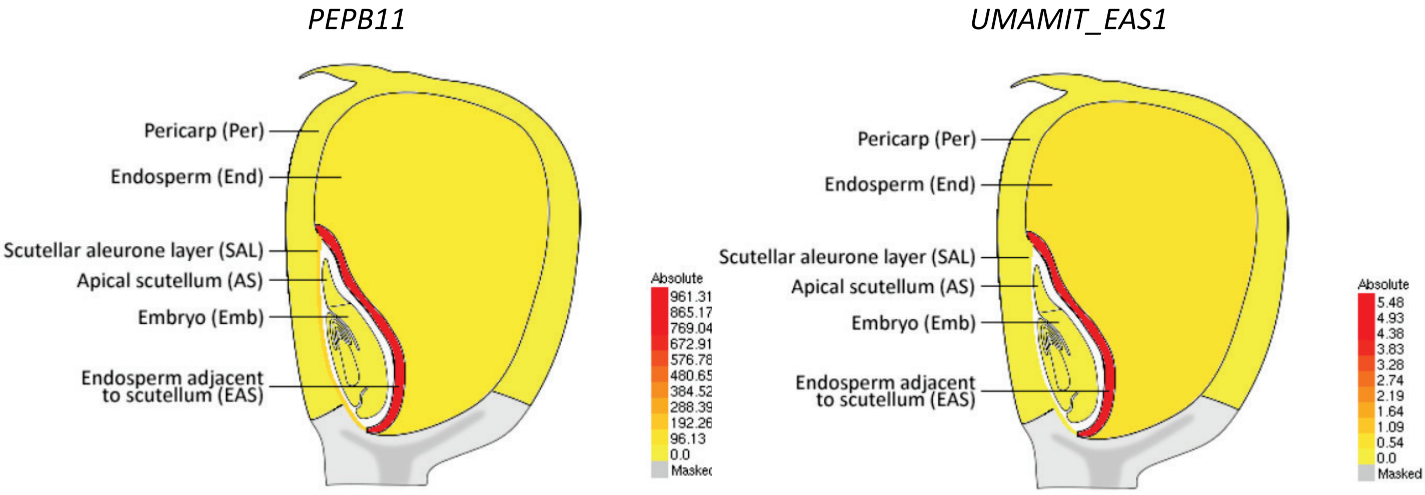


**Supplemental Figure 3.** Example of eFP Browser views (http://bar.utoronto.ca/efp_maize/cgi-bin/efpWeb.cgi?dataSource=Maize_Kernel) for two selected EAS marker genes used for *in-situ* hybridization experiment (Figure 3), and having the highest (*PEPB11*) and lowest (*UMAMIT_EAS1*) expression level in EAS (Supplemental Table 3). Note that the color scale reflecting genes expression level is automatically adjusted and could be manually configured using "Signal Threshold" option.

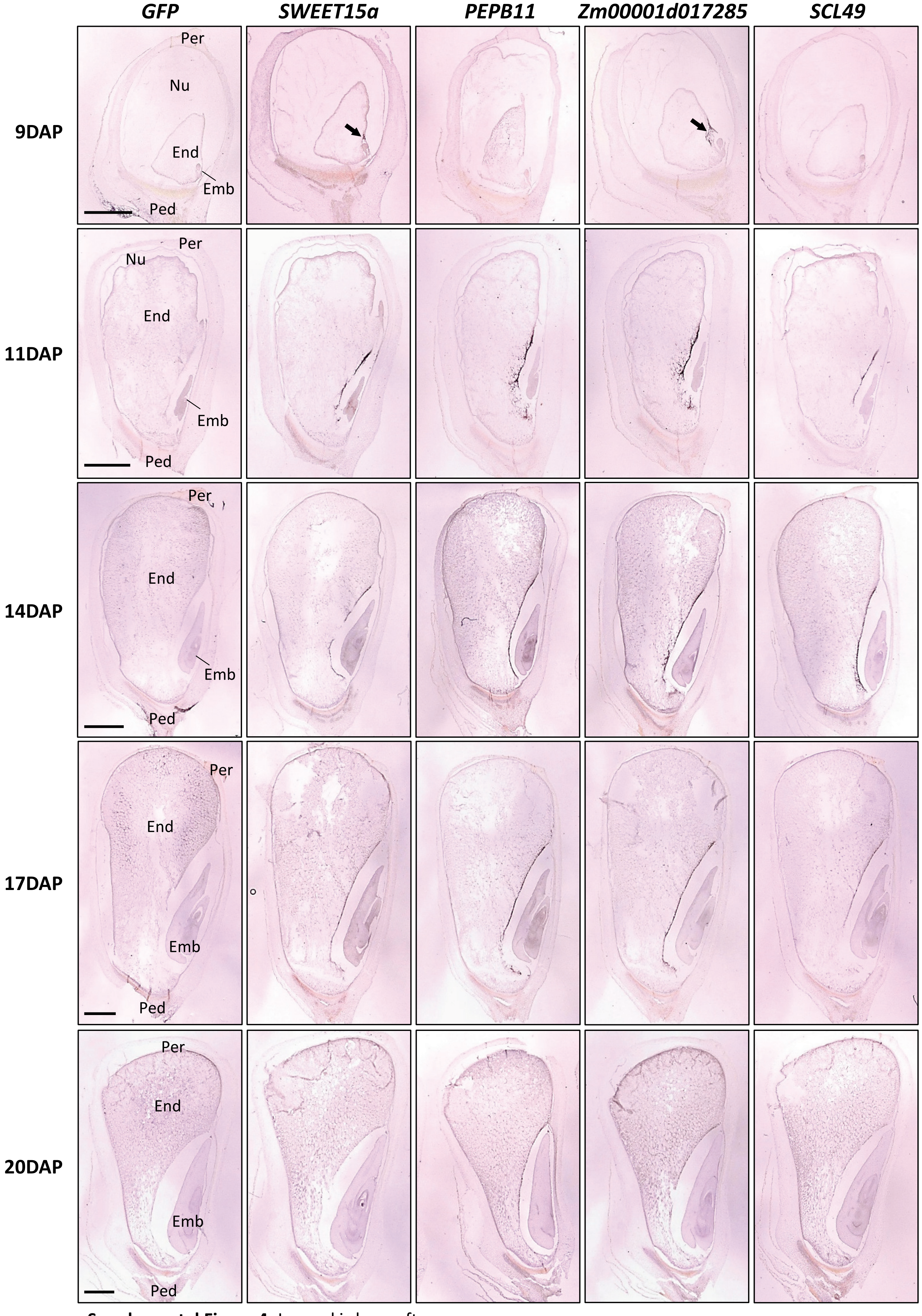


**Supplemental Figure 4.** Legend is here after.

**Supplemental Figure 4.** Pictures representing the the whole kernel for the *in situ* hybridizations presented in figure 4. Four probes detecting EAS marker genes (*SWEET15a*, *PEPB11*, *Zm00001d017285*, *SCL49*) were used on kernel sections at different stage. Scale bars corresponds to 1000 µm. For each photo the name of the probe is indicated at the top of the figure and the stage on the left. End = endosperm, emb = embryo, per = pericarp, nu = nucellus, ped = pedicel. Arrows indicate the main *in situ* hybridizations signal at 9DAP.

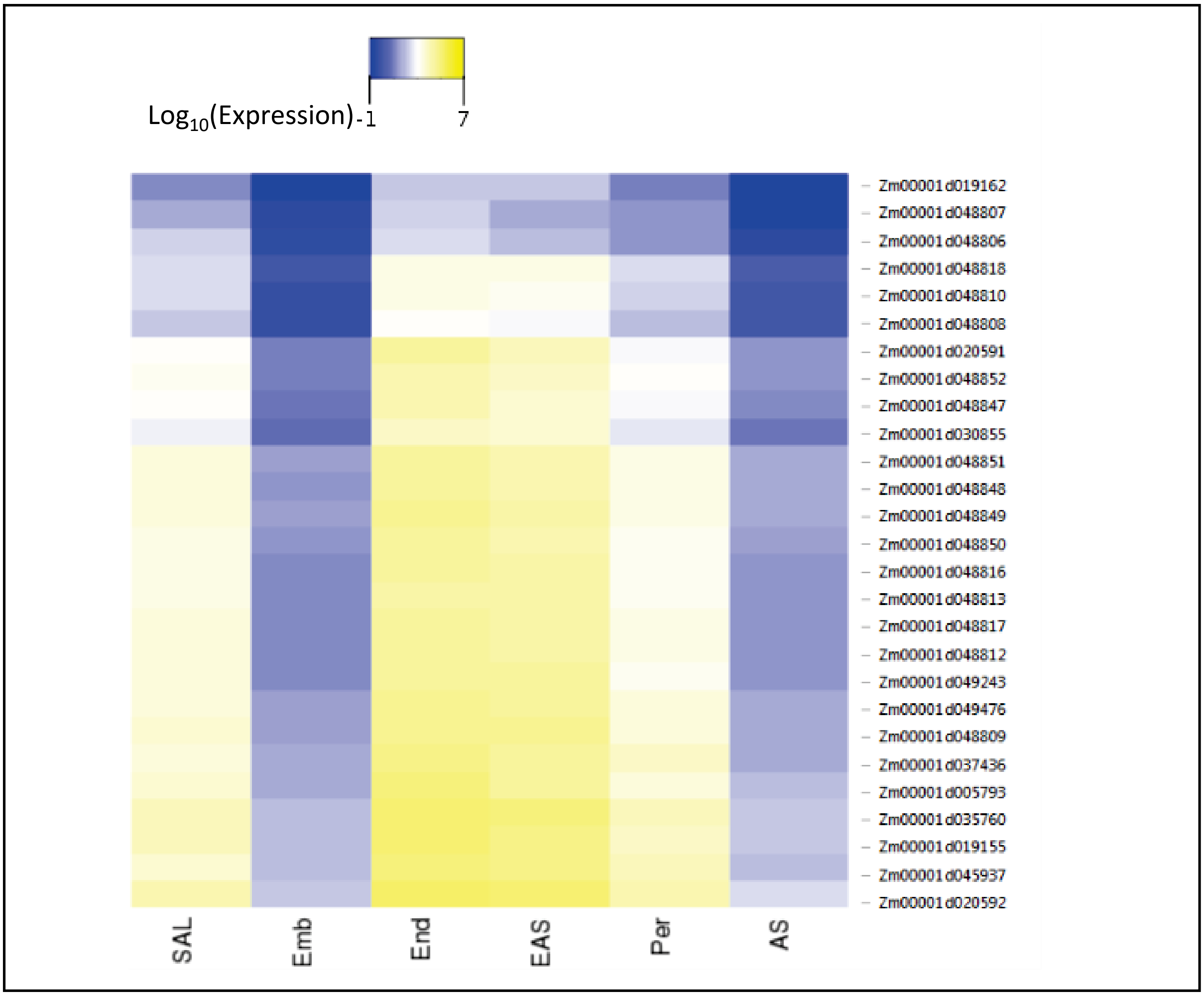


**Supplemental Figure 5.** Heat map of *Zein* precursor gene expression. For the (sub)compartments, the logarithm (base 10) of the average of the normalized read counts between the 4 replicates has been taken. An arbitrary value of -1 is given when the gene is not expressed in a (sub)compartment.

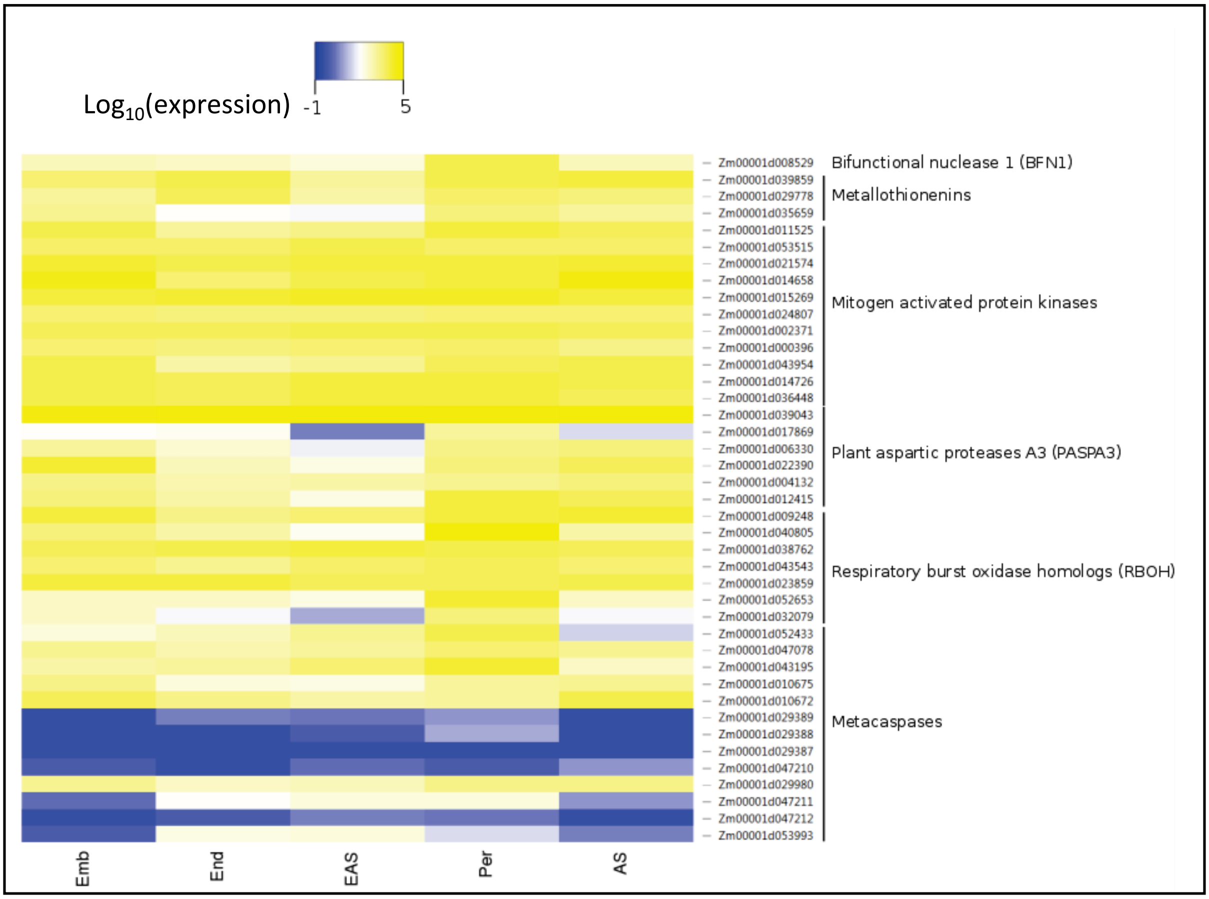


**Supplemental Figure 6.** Heat maps for genes potentially involved in programmed cell death. For the (sub)compartments, the logarithm (base 10) of the average of the normalized read counts between the 4 replicates has been taken. An arbitrary value of -1 is given when the gene is not expressed in a (sub)compartment.
